# Non-canonical glutamate-cysteine ligase activity protects against ferroptosis

**DOI:** 10.1101/2020.05.29.123802

**Authors:** Yun Pyo Kang, Andrea Mockabee-Macias, Chang Jiang, Isaac S. Harris, Gina M. DeNicola

## Abstract

Cysteine is required for maintaining cellular redox homeostasis in both normal and transformed cells. Deprivation of cysteine induces the iron-dependent form of cell death known as ferroptosis; however, the metabolic consequences of cysteine starvation beyond impairment of glutathione synthesis are uncharacterized. Here, we find that cystine starvation promotes ferroptosis not only through the inhibition of glutathione (GSH) synthesis, but also through the accumulation of glutamate. Surprisingly, we find that glutamate-cysteine ligase catalytic subunit (GCLC) prevents glutamate accumulation through the generation of alternative γ-glutamyl peptides. Further, inhibition of GCLC accelerates ferroptosis under cystine starvation in a GSH-independent manner. These results indicate that GCLC has an additional, non-canonical role in the protection against ferroptosis to maintain glutamate homeostasis under cystine starvation.

## Introduction

Amino acids can play critical biosynthetic functions beyond their use for protein synthesis. A notable example is the thiol-containing amino acid cysteine. Cysteine-derived molecules are crucial for multiple cellular processes as a consequence of their sulfur moiety, which facilitates diverse functions, including enzyme catalysis, energy transfer, and redox metabolism (Furuyama and Sassa, 2000; Martinez-Reyes et al., 2016; Rouault, 2012; Solmonson and DeBerardinis, 2018; Vyas et al., 2016). Cysteine is a rate-limiting substrate for the synthesis of glutathione (GSH) (Stipanuk et al., 2006), the most abundant intracellular antioxidant (Winterbourn and Hampton, 2008). GSH is a tripeptide consisting of the amino acids cysteine, glutamate and glycine. The synthesis of GSH occurs in two steps (Anderson, 1998). First, glutamate and cysteine are ligated by GCLC, producing the dipeptide γ-glutamyl cysteine (γ-Glu-Cys). Next, glycine is added to γ-Glu-Cys, producing the tripeptide GSH (γ-Glu-Cys-Gly). The antioxidant activity of GSH is a consequence of its function as a cofactor to multiple antioxidant proteins, including glutaredoxins (GRXs), GSH peroxidases (GPXs), and GSH S-transferases, thereby removing reactive oxygen species (ROS) (Harris and DeNicola, 2020).

Because of both its reactive thiol moiety and its essential function in redox homeostasis, cysteine levels are tightly regulated. While cysteine excess is prevented by overflow into the taurine pathway (Stipanuk et al., 2009), cysteine demand is met by inducible regulation of cystine import. Following oxidative stress, the expression of the cystine/glutamate exchange transporter xCT is induced, (Habib et al., 2015) thereby facilitating the uptake of cystine and its reduction to cysteine. In some tissues, most notably the liver, cysteine is also synthesized from homocysteine and serine via the transsulfuration pathway (Beatty and Reed, 1980; Rao et al., 1990; Reed and Orrenius, 1977). Given the important roles of cysteine, many cancers overexpress xCT (Ji et al., 2018; Takeuchi et al., 2013; Timmerman et al., 2013), which is positively regulated by oncogenic RAS (Lim et al., 2019) and NRF2 (Sasaki et al., 2002), and negatively regulated by the tumor suppressor p53 (Jiang et al., 2015). Pharmacological targeting of cystine uptake can effectively induce cancer cell death (Cramer et al., 2017; Dixon et al., 2012; Zhang et al., 2019), and cystine starvation can impair growth in multiple *in-vivo* cancer models (Cramer et al., 2017; Zhang et al., 2019).

Cysteine inadequacy can induce an iron-dependent form of cell death known as ferroptosis (Dixon et al., 2012). Ferroptosis is triggered by the reaction of polyunsaturated fatty acids (PUFA) in membrane lipids with peroxyl radicals produced from iron (Fe^2+^) and ROS (Cao and Dixon, 2016; Yang et al., 2014), thereby inducing lipid peroxidation. Consistently, processes that promote ferroptosis include increased ferritin uptake (Gao et al., 2015), ferritin degradation (Mancias et al., 2014), synthesis of PUFA containing lipids (Dixon et al., 2015; Doll et al., 2017), and mitochondrial ROS production (Gao et al., 2015; Gao et al., 2019). However, while cysteine is directly linked to GSH synthesis, which can influence the levels of ROS and lipid peroxides via GPX4 (Yang et al., 2014), cysteine availability can also influence the levels of cofactors and metabolites within cells beyond its use for GSH synthesis. Importantly, the metabolic consequences of cystine starvation are poorly understood.

To understand the metabolic consequences of the cystine starvation, we performed quantitative stable isotope labeled metabolite tracing in non-small cell lung cancer (NSCLC) cells. We found that glutamate accumulated due to impaired GSH synthesis, and promoted ferroptosis. Further, we identified that under cysteine-deprived conditions, GCLC used other small, non-charged amino acids in place of cysteine to generate γ-glutamyl peptides. This promiscuous activity prevented glutamate accumulation to protect against ferroptosis. γ-glutamyl peptide synthesis by GCLC was also evident in mouse tissues.

## Results

### Cystine starvation impairs GSH synthesis prior to the onset of ferroptosis

To evaluate the consequence of cystine starvation in NSCLC cells, we starved a panel of cell lines of extracellular cystine and first monitored viability over time. Cystine starvation induced the death of most cell lines between 24-48 hrs, with the exception of H460 and H1944 cells, which were more resistant (Figure 1A). Cell death was confirmed to be ferroptosis due to both the ability of the ferroptosis inhibitor Ferrostatin-1 (Fer-1) and iron chelator DFO (Dixon et al., 2012) to rescue cell death (Figure 1A) and the morphological changes characteristic of ferroptosis (Figure S1A). Next, we examined the metabolic consequences of cystine starvation. To understand the immediate consequences of cystine starvation, we starved cells for 4 hrs, which caused the rapid depletion of intracellular cysteine to almost undetectable levels in all cell lines (Figure 1B). To determine whether cystine starvation influenced cellular processes, we first examined glutathione (GSH) synthesis. Cysteine is a rate limiting metabolite of GSH synthesis (Stipanuk et al., 2006) and GSH also plays an important role in ferroptosis via ROS metabolism (Dixon et al., 2012) and substrate of GPX4 (Conrad and Friedmann Angeli, 2015). Therefore, to evaluate the effect of extracellular cystine starvation on GSH synthesis, we conducted quantitative ^13^C_3_-serine tracing. ^13^C_3_-serine can be metabolized to ^13^C_2_-glycine (M+2) and ^13^C_3_-cysteine (M+3), which are subsequently incorporated into GSH (M+2 and M+3, respectively, Figure 1C). After 4 hours labeling, most of the serine fraction and half of the glycine fraction were labeled (Figure S1B and S1C). Importantly, while the amount of newly synthesized glycine was equivalent or increased in the cell lines following cystine starvation, the amount of M+2 glycine incorporated into GSH was dramatically depleted. Minimal M+3 labeling was detected. In addition, total GSH levels were lower, consistent with an inhibition of GSH synthesis. These results indicate that the extracellular cystine starvation rapidly depletes intracellular cysteine availability for GSH synthesis, which precedes the induction of ferroptosis.

**Figure 1.**
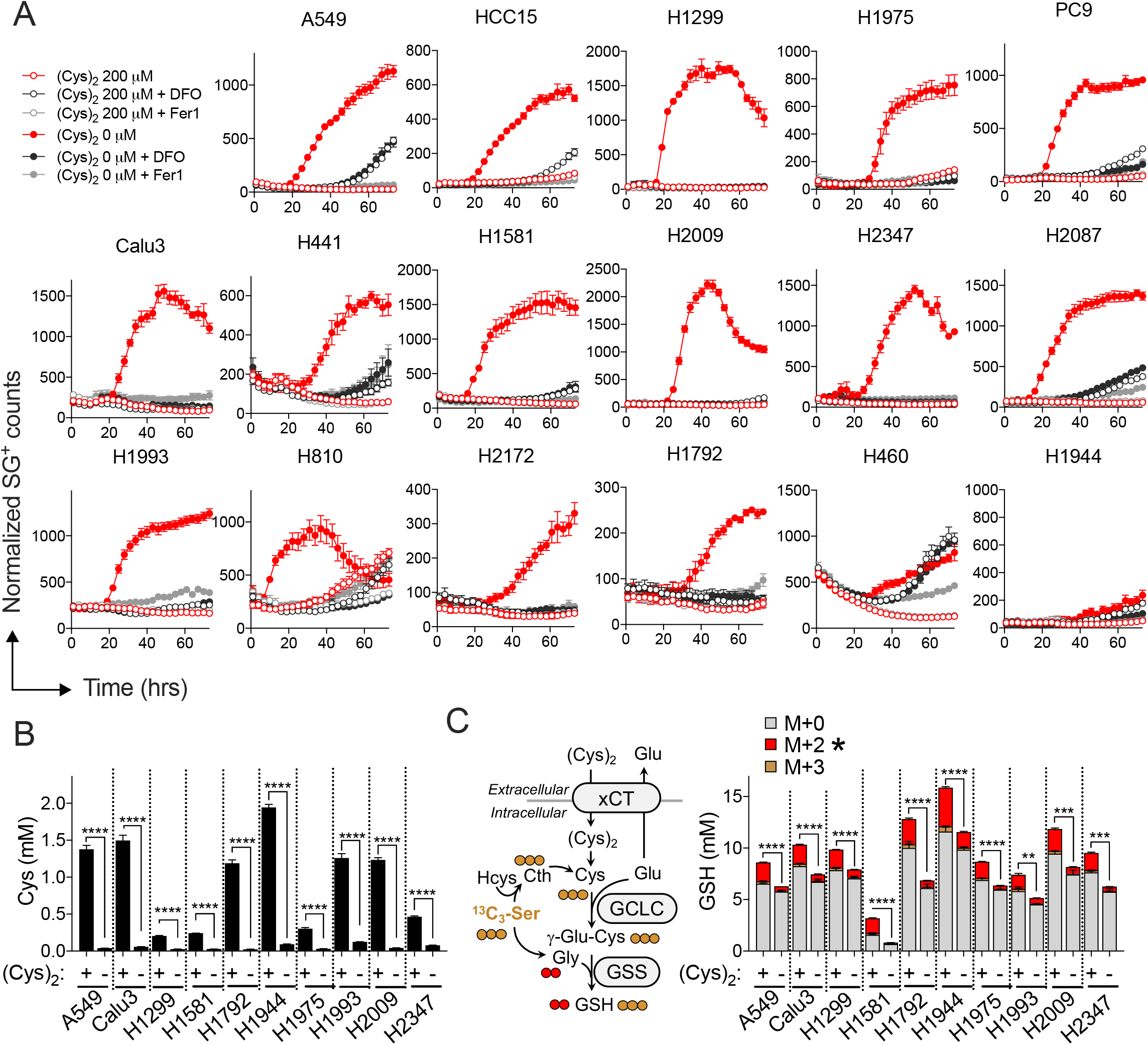
Cysteine starvation induces ferroptosis and impairs GSH synthesis. (A) Time course Measurement of NSCLC cell death under cystine starved (0μM) or replete (200μM) conditions treated with Vehicle (0.1% DMSO), Ferrostatin-1 (Fer-1, 10 μM) or DFO (100 μM). (N=4). Cell death was determined by Sytox Green staining and normalized to cell density. (B) Quantitation of intracellular cysteine (N=3) and (C) quantitative [^13^C_3_]-serine tracing into glutathione following media change to cystine starved (-) or replete (+) conditions for 4 hrs. (N=3) Data shown as mean ± SEM. N is number of biological replicates. **P<0.01, ***P<0.001, and ****P<0.0001. Unpaired two-tailed t test between M+2 labeling fractions was used for the statistical comparisons in (B).

### Inhibition of GSH synthesis promotes glutamate accumulation

GSH synthesis consumes glutamate and glycine in addition to cysteine. We observed that inhibition of GSH synthesis was associated with an accumulation of intracellular glycine and glutamate in multiple NSCLC cell lines following cystine starvation (Figure 2A). In addition, glutamate export is obligatory for cystine import and glutamate accumulation may also be influenced by reduced cystine/glutamate exchange. Consistently, we observed that cystine starvation decreased glutamate exportation (Figure 2A and D). Because glutamate was previously shown to contribute ferroptosis (Gao et al., 2015), we examined whether glutamate plays a causal role in cystine-starvation induced ferroptosis in NSCLC cells. We treated cells with 5 mM glutamate diethyl ester (GlutEE), a concentration we confirmed increases intracellular glutamate to similar levels as cystine starvation in A549 cells (Figure S2A and 2A). We found that GlutEE treatment accelerated ferroptosis in multiple NSCLC cells (Figure 2B), while glutamine starvation, which depleted intracellular glutamate (Figure S2D), or treatment with the transaminase inhibitor AOA could rescue ferroptosis (Figure 2C). Interestingly, the effects of AOA could be overridden by treatment with dimethyl-alpha-ketoglurate (DMαKG), suggesting αKG or its downstream metabolite mediates the effects of glutamate. Glutamate was previously shown to promote ferroptosis via the TCA cycle and ROS generated from the oxidative phosphorylation (Gao et al., 2015; Gao et al., 2019). Consistently, we found that GlutEE promoted ROS accumulation under cysteine starved conditions (Figure S2B). Therefore, these results demonstrate that inhibition of GSH synthesis by cystine starvation not only depletes GSH, but also induces the accumulation of glutamate to promote ferroptosis.

**Figure 2.**
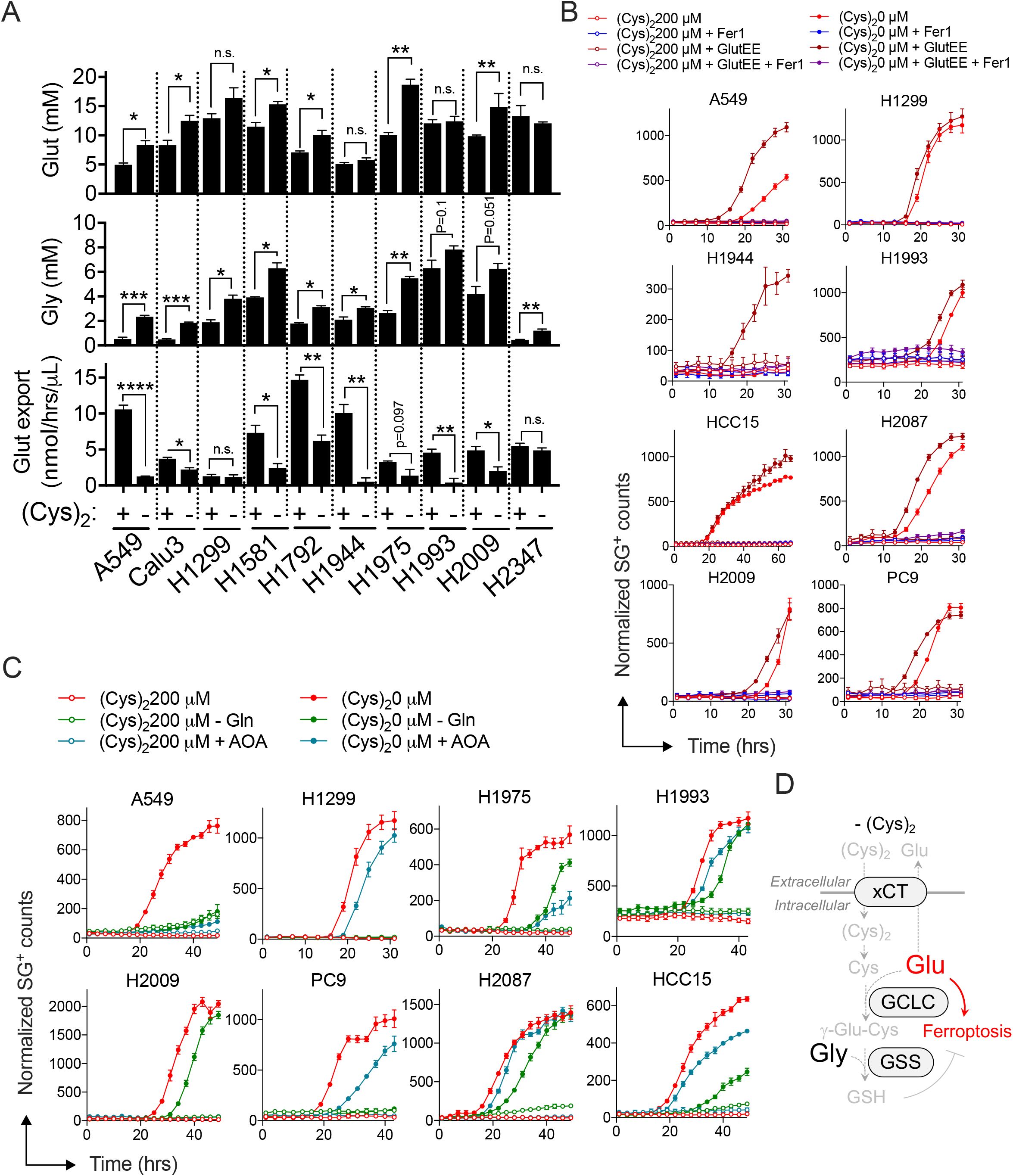
Inhibition of GSH synthesis promotes ferroptosis via glutamate accumulation. (A) Quantitation of intracellular glutamate (Glut, upper) and glycine (Gly, middle), and exportation rate of glutamate (bottom) in NSCLC cell lines under cystine replete or starved conditions for 12 hrs. (N=3). (B) Measurement of NSCLC cell death under cystine starved or replete conditions treated with Vehicle, Fer-1 (10 uM), and/or glutamate diethyl ester (GlutEE, 5 mM) as indicated (N=4 except H1299, cystine 0 μM + GlutEE group: N=3). (C) Measurement of NSCLC cell death under cystine starved or replete conditions treated with Vehicle or AOA (0.5 mM), or without media glutamine (-Gln) (N=4). Vehicle-treated cystine replete and starved data are from (B). For A-C, data are shown as mean ± SEM (A, B, and C). n.s., not significant; *P<0.05, **P<0.01, ***P<0.001, and ****P<0.0001. N is number of biological replicates. An unpaired two-tailed t test was used for statistical analysis in (A). (D) Schematic depiction of glutamate accumulation-mediated ferroptosis promotion under cystine starved conditions.

### GCLC prevents ferroptosis independent of GSH production

Multiple studies have demonstrated a potent, synergistic effect of GSH synthesis inhibition with BSO with limitation of cystine uptake or availability (Cramer et al., 2017; Harris et al., 2015). Consistently, we observe that BSO treatment promotes ferroptosis under cystine starvation in most NSCLC cell lines (Figure 3A). However, we find that cystine starvation rapidly inhibits intracellular GSH synthesis (Figure 1B). Importantly, while BSO treatment depleted GSH in cystine replete cells as expected, it did not change GSH levels in cystine starved cells (Figure 3B). Thus, the accelerated cell death induced by BSO under cystine starvation may not be explained by the depletion of GSH. Therefore, we examined whether GCLC could play a role in cystine-starvation induced ferroptosis independent of GSH synthesis. To evaluate this question, we generated GCLC and GSS KO A549 clones (Figure 3C). Importantly, both clones were defective in GSH synthesis as evidenced by significantly reduced intracellular GSH levels effectively compared to parental cells (Figure 3D). Importantly, the GCLC KO clones demonstrated accelerated ferroptosis induction under cystine starvation compared to parental cells, which could be rescued by GCLC cDNA, while the GSS KO clones did not (Figure 3E). Further, BSO treatment induced ferroptosis in the GSSKO clone (Figure 3F), despite the absence of GSH (Figure 3D), further confirming the GSH-independent role of GCLC in ferroptosis protection. Finally, GCLC KO, but not GSS KO, accelerated ferroptosis induction under cystine starvation following acute deletion in H1299 cells (Figure S3A-C), suggesting this is not a consequence of adaptation in single cell clones. Together, these data indicate that GCLC can prevent cystine starvation-induced ferroptosis of NSCLC cells independent of GSH production (Figure 3G).

**Figure 3.**
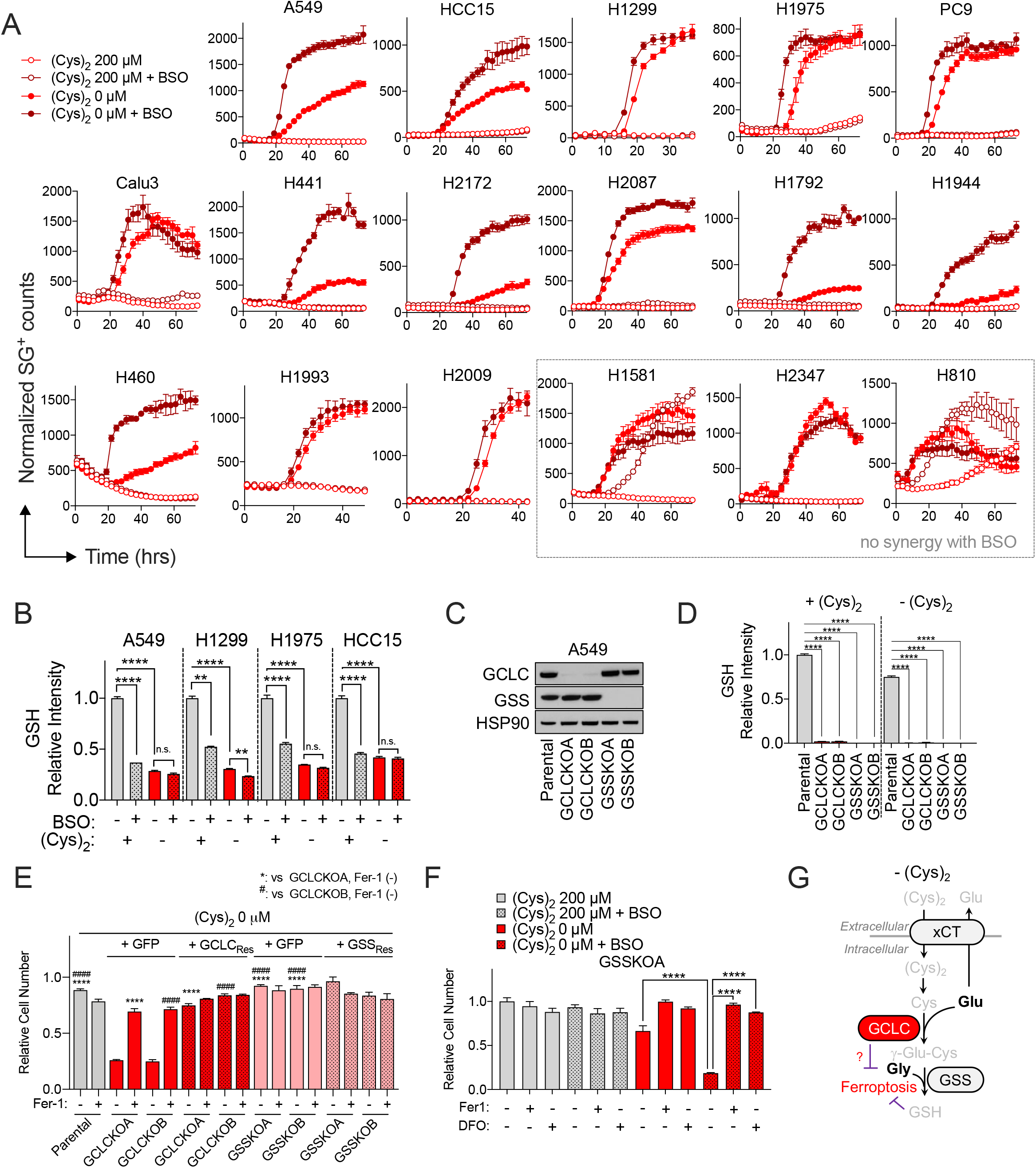
GCLC prevents ferroptosis independent of GSH production. (A) Measurement of NSCLC cell death under cystine starved or replete conditions treated with Vehicle (0.1% DMSO) or BSO (100 μM). (N=4). Vehicle-treated cystine replete and starved data are from Fig. 1A. (B) Intracellular glutathione levels under cystine starved or replete conditions treated with Vehicle (0.1% DMSO) or BSO (100 μM) for 13 hrs. (N=3). Data was normalized by the mean value of vehicle-treated cystine replete conditions. (C) Representative immunoblots of GCLC, GSS, and HSP90 (loading control) from A549 GCLC KO clones (GCLCKOA and GCLCKOB), GSS KO clones (GSSKOA and GSSKOB), and parental A549 cells. (D) Intracellular GSH levels of A549 GCLC KO clones, GSS KO clones, and parental cells under cystine replete or starved conditions for 3 hrs (N=3). Data was normalized by mean value of parental cells under cystine replete conditions. (E) Relative cell number of parental A549 cells, GCLC KO clones reconstituted with GFP (+GFP) or sgRNA-resistant GCLC (+GCLC_Res_), and GSS KO clones reconstituted with GFP (+GFP) or sgRNA-resistant GSS (+GSS_Res_) under cystine starved condition treated with vehicle or Fer-1 (10 μM) as indicated for 16 hrs (N=3). Data are normalized to the mean value of vehicle-treated cystine replete conditions. (F) Relative cell number of A549 GSSKO cells under cystine starved conditions treated with vehicle (0.1% DMSO), Fer-1 (10 μM), or DFO (100 μM) as indicated for 16 hrs (N=3). Data are normalized to the mean value of vehicle-treated cystine replete conditions. For A, B, D-F, data are shown as mean ± SEM. N is number of biological replicates. n.s., not significant; **P<0.01, ****P<0.0001, and ^####^P<0.0001. For B, D, E, F, a oneway ANOVA with Bonferroni’s multiple comparison test was used for statistical analyses. (G) Schematic depicting the glutathione synthesis-independent role of GCLC to prevent ferroptosis under cystine starved conditions.

### Cystine starvation induces GCLC-dependent γ-glutamyl peptide accumulation

To determine the mechanism of GSH-independent protection against ferroptosis by GCLC, we conducted non-targeted metabolomics. Interestingly, we discovered a cluster of LC-MS peaks which were highly depleted by BSO treatment following cystine starvation in A549 cells (Figure 4A). Further, these peaks were the same ones that were the most highly accumulated by extracellular cystine starvation (Figure 4A). These unknown LC-MS peaks were identified as γ-glutamyl-di or tri-peptides, which all contain a glutamate-derived moiety (Figure 4A). Authentic standards for γ-glutamyl threonine (γ-Glu-Thr) and γ-glutamyl-alanyl-glycine (γ-Glu-Ala-Gly) were not available, thus we further validated their identity via stable isotope labeled metabolite tracing. The ^13^C_5_, ^15^N_2_-glutamine tracing result indicated that both γ-Glu-Thr and γ-Glu-Ala-Gly were derived from glutamate (Figure 4D). Further, 2, 3, 3-^2^H_3_-serine tracing showed that γ-Glu-Ala-Gly was derived from the glycine (Figure S4A). We extended these observations to other NSCLC cell lines and found that cystine starvation consistently promoted the accumulation of γ-glutamyl peptides, which was inhibited by treatment with BSO (Figure 4B). These results suggest that cystine starvation promotes the accumulation of γ-glutamyl peptides by the GSH synthesis pathway.

**Figure 4.**
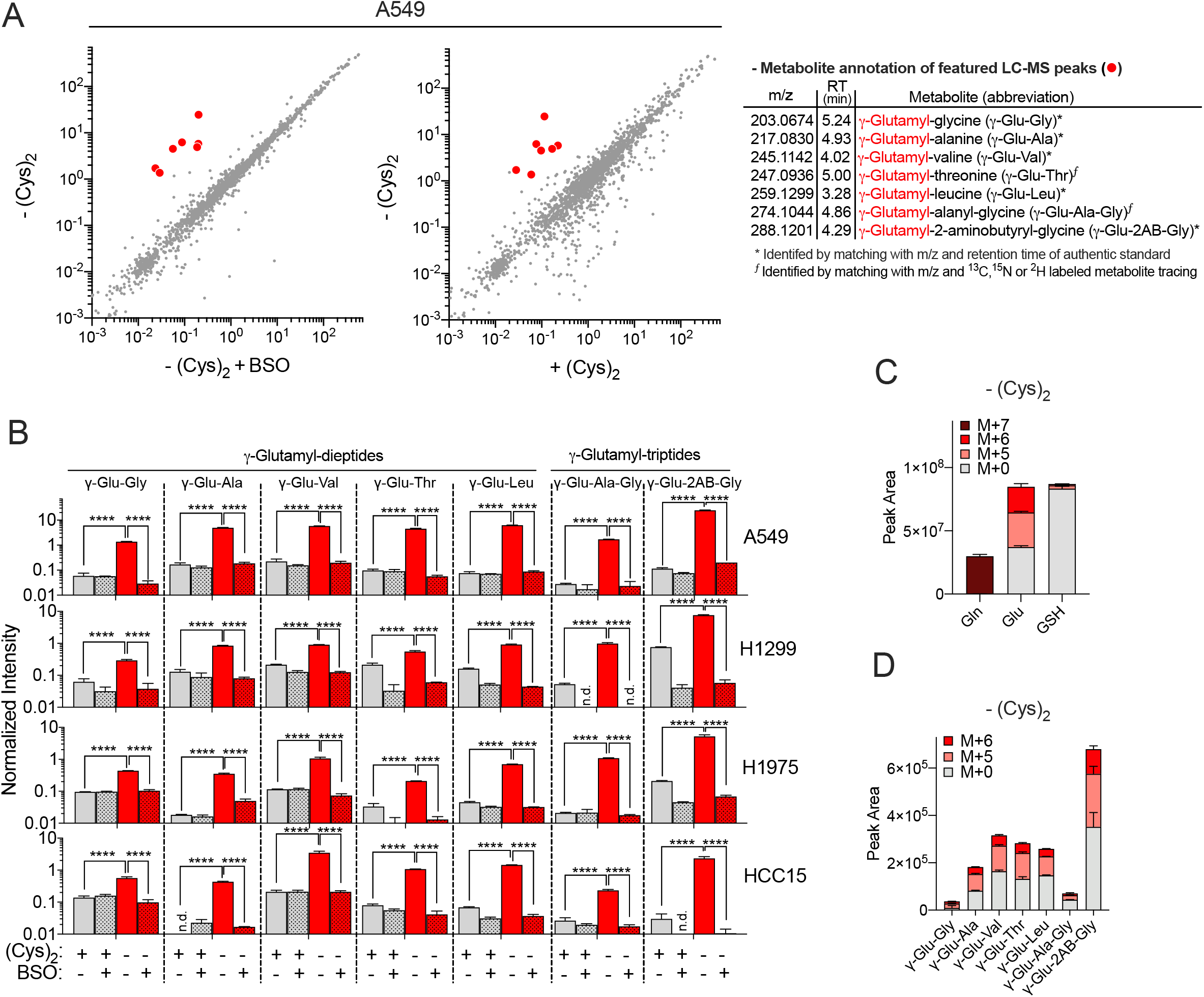
Cystine starvation promotes γ-Glut-dipeptide production by GCLC. (A) Scatter plot comparison of non-targeted metabolomics features in A549 cells (Left) between control and BSO treated groups under cystine starved conditions and (middle) between cystine replete and starved conditions. The mean intensity of median-normalized LC-MS peaks of each group (N=3) are plotted on the axes, and each dot represents an individual LC-MS peak. The highly altered LC-MS peaks (red dots) were further identified and annotated (right). (B) The intensity of γ-glutamyl di- and tri-peptides in 4 NSCLC cells following cystine replete or starved condition and treatment with vehicle or without BSO (100 μM) for 13 hrs (N=3). Data are normalized to the median value of all LC-MS features in each sample. ^13^C_5_, ^15^N_2_-Gln tracing of A549 cells into (C) Gln, Glut, GSH, and (D) γ-Glut-peptides following cystine starvation for 4 hrs (N=3). For B-D, data are shown as mean ± SEM. N is number of biological replicates. n.d., not detected; ****P<0.0001. For B, a one-way ANOVA with Bonferroni’s multiple comparison test was used.

### γ-glutamyl peptide synthesis by GCLC scavenges glutamate to protect against ferroptosis

The tripeptide γ-glutamyl-2-aminobutyryl-glycine (γ-Glu-2AB-Gly) is known to be generated by GCLC and GSS (Huang et al., 1988; Oppenheimer et al., 1979) in a similar manner to GSH by substituting 2-aminobutyrate for cysteine (Figure S4B). Consequently, the accumulation of γ-Glu-2AB-Gly under cysteine starvation can be explained by cysteine unavailability for GCLC (Figure 4B and S4B). In contrast, γ-glutamyl-dipeptides are reported to be derived from GSH by γ-glutamyl transferase (GGT) extracellularly (Figure S4B) (Hanigan and Pitot, 1985). However, ^13^C_5_, ^15^N_2_-glutamine tracing demonstrated that while the newly labeled GSH fraction was very small, as expected, glutamate and γ-glutamyl-dipeptides were newly labeled to ~ 50% in cystine starved A549 cells (Figure 4C and D), suggesting that the γ-glutamyl dipeptides were synthesized from glutamate but not from GSH (Figure S4B). Because γ-glutamyl-valine is synthesized by the *Saccharomyces cerevisiae* glutamate-cysteine ligase (Sofyanovich et al., 2019), and γ-glutamyl dipeptide synthesis by mouse liver extracts was recently shown to be GCLC-dependent (Kobayashi et al., 2020), we hypothesized that the γ-glutamyl dipeptides were directly generated by GCLC rather than GSH metabolism by GGT. To evaluate this, we evaluated the γ-glutamyl dipeptide levels in the GCLC and GSS KO clones. Importantly, the γ-glutamyl dipeptides that were accumulated following cystine starvation in parental cells were dramatically depleted only in the GCLC KO clones (Figure 5A). Interestingly, the γ-glutamyl dipeptide levels were generally higher in GSS KO clones than parental lines, which can be explained by the feedback inhibition of GCLC by GSH, and more weakly by γ-Glu-2AB-Gly (Richman and Meister, 1975) (Figure 5A). In addition, both γ-Glu-2AB-Gly and γ-Glu-Ala-Gly tripeptides were depleted by both GCLC and GSS KO compared to parental cells, as GSS activity is required for the ligation of glycine. Consistent alterations of γ-glutamyl peptides were observed in GCLC and GSS KO H1299 cells, which were rescued by GCLC or GSS restoration (Figure S5A). These results indicate that γ-glutamyl dipeptides are directly generated by GCLC under cystine starved condition (Figure 5B).

**Figure 5.**
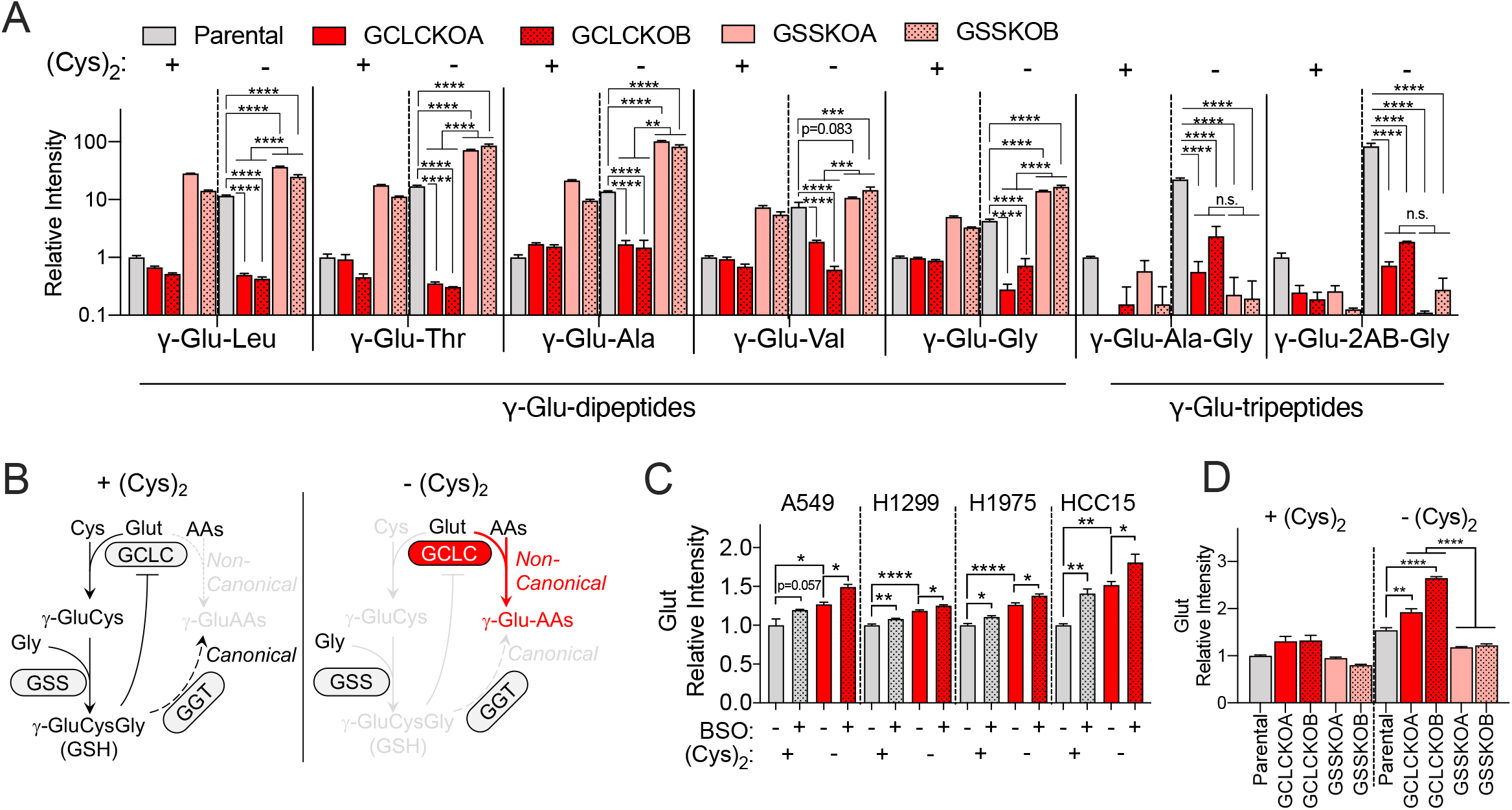
γ-glutamyl peptide synthesis by GCLC scavenges glutamate to protect against ferroptosis. (A) Intracellular γ-glutamyl peptide levels in parental A549s and GCLC and GSS KO clones following cystine replete or starved conditions for 3 hrs (N=3). Data was normalized by the mean value of parental lines under cystine replete conditions. (B) Schematic depicting the non-canonical, γ-glutamyl dipeptide synthesis function of GCLC under cystine starvation. GCLC consumes glutamate to produce alternative γ-glutamyl amino acids. (C) Intracellular glutamate levels following cystine replete or starved conditions and treatment with vehicle or BSO (100 μM) for 13 hrs (N=3). Data were normalized to the mean value of vehicle-treated cystine replete conditions. (D) Intracellular glutamate levels in parental A549 cells and GCLC and GSS KO clones following cystine replete or starved condition for 3 hrs (N=3). Data are normalized to the mean value of cysteine-replete parental lines. For A, C, and D, data are shown as mean ± SEM. N is number of biological replicates. n.d., not detected; n.s., not significant; *P<0.05, **P<0.01, ***P<0.001, and ****P<0.0001. For A, C, and D, a one-way ANOVA with Bonferroni’s multiple comparison test was used.

As we found that GCLC inhibition with BSO treatment or genetic KO could accelerate ferroptosis under cystine starvation (Figure 3A and E), we examined whether GCLC-mediated γ-glutamyl dipeptide synthesis plays a causal role in this process. Because glutamate accumulation promoted ferroptosis (Figure 2), we evaluated the ability of γ-glutamyl dipeptides to serve as a glutamate sink. Both inhibition of GCLC with BSO treatment and GCLC KO increased intracellular glutamate levels under cystine starvation, while GSS KO was actually protective (Figure 5C and D). We also observed an accumulation of glutamate in GCLC KO, but not GSS KO H1299 cells under cystine starvation (Figure S5B). Importantly, the γ-glutamyl dipeptides themselves did not play a protective role against ferroptosis as their supplementation did not rescue cystine starvation-induced ferroptosis of GCLC KO clones (Figure S5C). Finally, cystine starvation-induced ferroptosis of GCLC KO cells was rescued by both glutamine starvation and AOA treatment (Figure S5D). Together, these results demonstrate that GCLC has a non-canonical role in ferroptosis to balance the glutamate pool to protect against ferroptosis under cystine starvation (Figure S5E).

### GCLC regulates glutamate homeostasis in vivo

Finally, we examined whether GCLC mediates the synthesis of γ-glutamyl peptides in vivo under normal physiological conditions. To this end, systemic Gclc deletion was induced in an adult mouse (Figure 6A) and we examined the effect in the liver, kidney and serum. Efficacy of Gclc deletion was evident by the depletion of glutathione by 75-90% in these tissues (Figures 6B-D). While glutathione was present in the reduced (GSH) form in tissues, the serum had predominantly the oxidized form (GSSG), which may either be due to the oxidizing extracellular conditions or oxidation during sample preparation. Further, we found that Gclc KO liver, kidney, and serum were also depleted of γ-glutamyl-peptides, including both the dipeptides and tripeptides (Figures 6B-D). In addition, deletion of Gclc increased glutamate levels in the liver and serum, but not the kidney (Figures 6B-D). Overall, these results indicate that GCLC plays a causal role in the homeostatic control of glutamate and γ-glutamyl peptide metabolism in vivo (Figure 6E).

**Figure 6.**
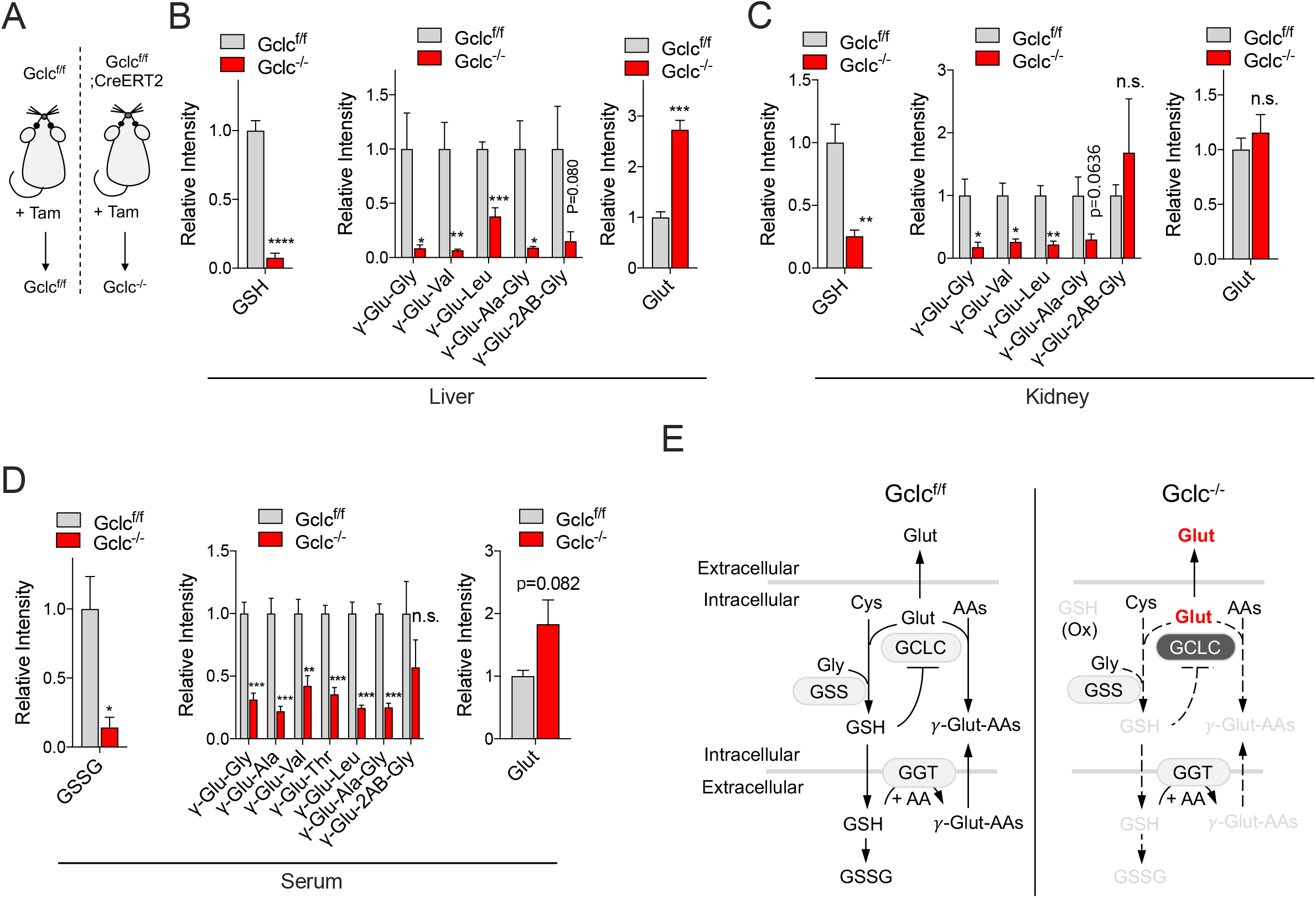
GCLC regulates glutamate homeostasis in vivo. (A) Tamoxifen (Tam)-inducible deletion in the adult mouse. Gclc^f/f^ (control, Gclc functional) and Gclc^-/-^ (Gclc knockout). Measurement of GSH, γ-Glu-peptides, and Glut levels in (B) liver and (C) kidney of Gclc^f/f^ and Gclc^-/-^ mice. Measurement of GSSG, γ-Glu-peptides, and Glut levels of (D) serum in Gclc^f/f^ and Gclc^-/-^ mice. For B-D, Data were normalized to the mean value of Gclc^f/f^ group. For B-D, data are shown as mean ± SEM. N is number of biological replicates. n.s., not significant; *P<0.05, **P<0.01, ***P<0.001, and ****P<0.0001. For B-D, Unpaired two-tailed t test was used for the statistical comparisons. (E) Schematic depiction of GCLC-mediated metabolic alterations in mouse tissue.

## Discussion

The findings reported herein demonstrate that γ-glutamyl peptide synthesis by GCLC provides GSH-independent protection from ferroptosis following cystine starvation. While cystine starvation-induced ferroptosis has commonly been attributed to the depletion of cellular GSH, we show that cystine starvation induces complex metabolic changes within cells. Our work does not exclude the importance of GSH in the protection against ferroptosis. GSH is a major intracellular antioxidant and substrate for GPX4 for lipid hydroperoxide detoxification (Dixon and Stockwell, 2019; Yang et al., 2014). However, our work demonstrates that inhibition of GSH synthesis causes a metabolic imbalance and accumulation of the amino acids glycine and glutamate, which plays a causal role in ferroptosis induction. Our findings are consistent with prior reports demonstrating that glutamate contributes to ferroptosis via ROS generation in the mitochondria (Gao et al., 2015; Gao et al., 2019). Previous studies have found that combined inhibition of cystine uptake with glutathione synthesis can synergistically inhibit the viability of cells and tumors (Cramer et al., 2017; Harris et al., 2015). Our work has important implications for the interpretation of studies using BSO to inhibit GCLC. While many of those results may be attributed to GSH depletion, the contribution of GCLC to γ-glutamyl peptide synthesis and glutamate scavenging may also play a very important role, particularly in the context of xCT inhibition, where cells cannot export glutamate. Our findings warrant the development of potent GSS inhibitors for the study of ferroptosis to distinguish these mechanisms. These inhibitors would also be valuable for therapeutic combinations with ferroptosis inducers, although they may increase glutamate scavenging, which may affect cellular responses.

Our in vivo results provide direct genetic evidence to support the GCLC-mediated γ-glutamyl peptide production that was recently been reported in mouse liver extracts (Kobayashi et al., 2020), where glutamate could be ligated with other amino acids in a reaction inhibited by BSO. The promiscuity of GCLC toward amino acids other than cysteine is not a unique feature of this enzyme, and has been shown for many other metabolic enzymes. For example, serine palmitoyl transferase will also metabolize alanine or glycine when serine is limiting (Penno et al., 2010) and glutamate-aspartate aminotransferase will also metabolize cysteine sulfinic acid (Weinstein et al., 1988). In the case of GCLC, this feature may have been selected for during evolution, as the *S. cerevisiae* homolog (Gsh1p) also has the ability to at least use valine (Sofyanovich et al., 2019). Additional work is needed to determine which other amino acids are accepted by *S. cerevisiae* Gsh1p. For the human enzyme, small, non-charged amino acids that are structurally similar to cysteine can be used based on their appearance in γ-glutamyl peptides, although the full spectrum of amino acids has not been tested in a direct enzymatic assay.

Our findings may extend beyond conditions of cysteine deficiency. Systemic deletion of mouse *Gclc* revealed that Gclc plays a role in the regulation of glutamate and γ-glutamyl peptides levels in normal tissue. Notably, glutamate accumulation was only observed in the liver but not the kidney. Liver plays a critical role in GSH synthesis to supply the rest of the organism, which may consume significantly more glutamate in liver than kidney (Ookhtens and Kaplowitz, 1998). Similarly, cancer cells synthesize a significant amount of GSH (Balendiran et al., 2004; Huang et al., 2001; Soini et al., 2001; Sun et al., 2019; Tatebe et al., 2002) and use glutamate for cystine export (Ji et al., 2018; Shin et al., 2017; Takeuchi et al., 2013; Timmerman et al., 2013), which may explain the robust accumulation of glutamate following cystine starvation in our NSCLC cells. It is important to note that, in contrast to cell culture, depletion of γ-glutamyl peptides in Gclc KO tissue may be a consequence of both canonical, extracellular GGT mediated γ-glutamyl dipeptide production and the intracellular GCLC-mediated pathway. However, the accumulation of glutamate and the depletion of γ-glutamyl tripeptides, which require the activity of GSS, strongly suggests that these peptides are being produced intracellularly. Supportively, the activity of GGT is negligible in the mouse liver compared to kidney (Kobayashi et al., 2020). It is not known whether γ-glutamyl peptides have additional functions in tissues beyond serving as a reservoir for glutamate, and potentially other amino acids. γ-glutamyl peptides levels have been shown to be increased under conditions of liver injury, including drug-induced injury, hepatitis infection, liver cirrhosis, and hepatocellular carcinoma (Soga et al., 2011). Additional work is needed to understand the role of γ-glutamyl peptide synthesis in these diseases.

We also find that GSS may regulate glycine homeostasis by producing γ-glutamyl tripeptides, including γ-Glu-2AB-Gly and γ-Glu-Ala-Gly. Future work is needed to both understand whether GSS can use other amino acids besides glycine and determine the full spectrum of γ-glutamyl tripeptides produced by GSS. Further, we find that GSS deficiency actually enhances γ-glutamyl dipeptide synthesis, which can be explained by loss of feedback inhibition of GCLC by GSH. These findings raise interesting implications for the metabolic phenotypes of patients with inborn errors of glutathione metabolism. Although extremely rare, mutations in GCLC and GSS result in hemolytic anemia. Interestingly, GSS mutant patients also present with 5-oxo-prolinuria, which is not observed in GCLC deficiency (Ristoff and Larsson, 2007). While this 5-oxo-prolinuria has been attributed to the accumulation of γ-glutamylcysteine and its metabolism to 5-oxo-proline by γ-glutamylcyclotransferase (GGCT) (Ristoff and Larsson, 2007), our work suggests that other γ-glutamyl-amino acids are likely produced in this situation to contribute to 5-oxo-prolinuria. This is likely to depend on the availability of cysteine, which would likely become limiting if the feedback inhibition of GCLC is lost due to an inability to synthesize GSH.

## Key Resources

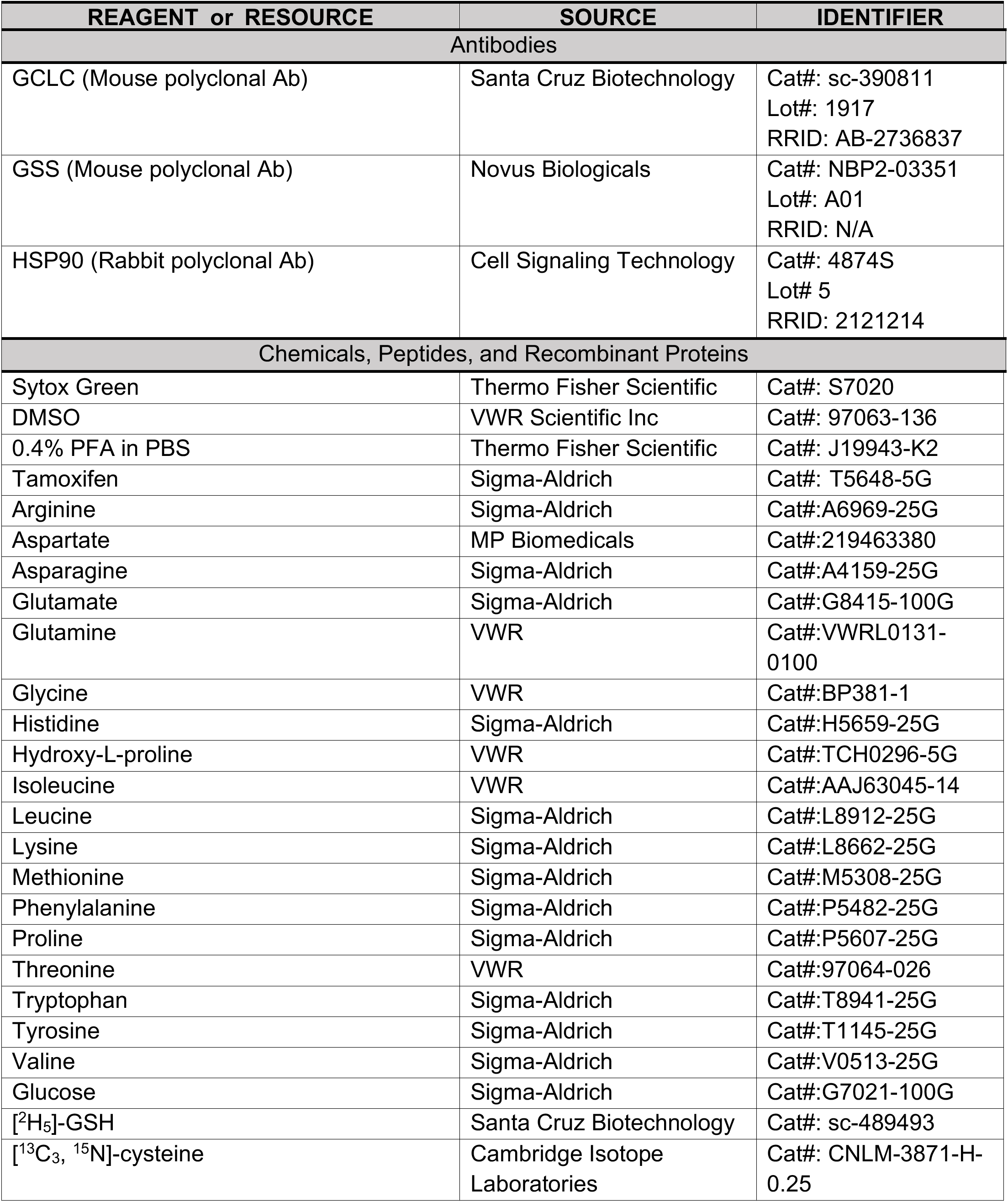
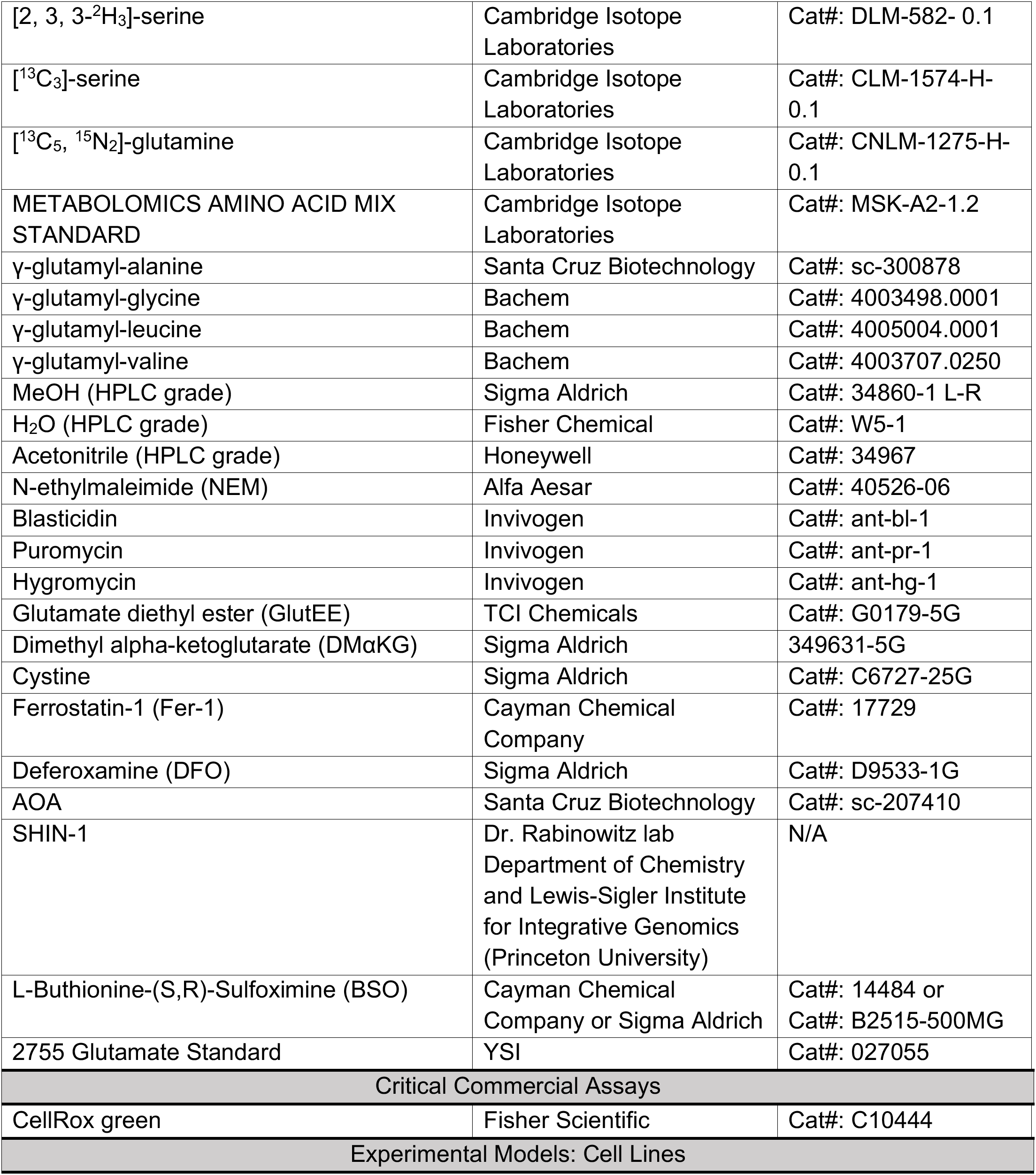
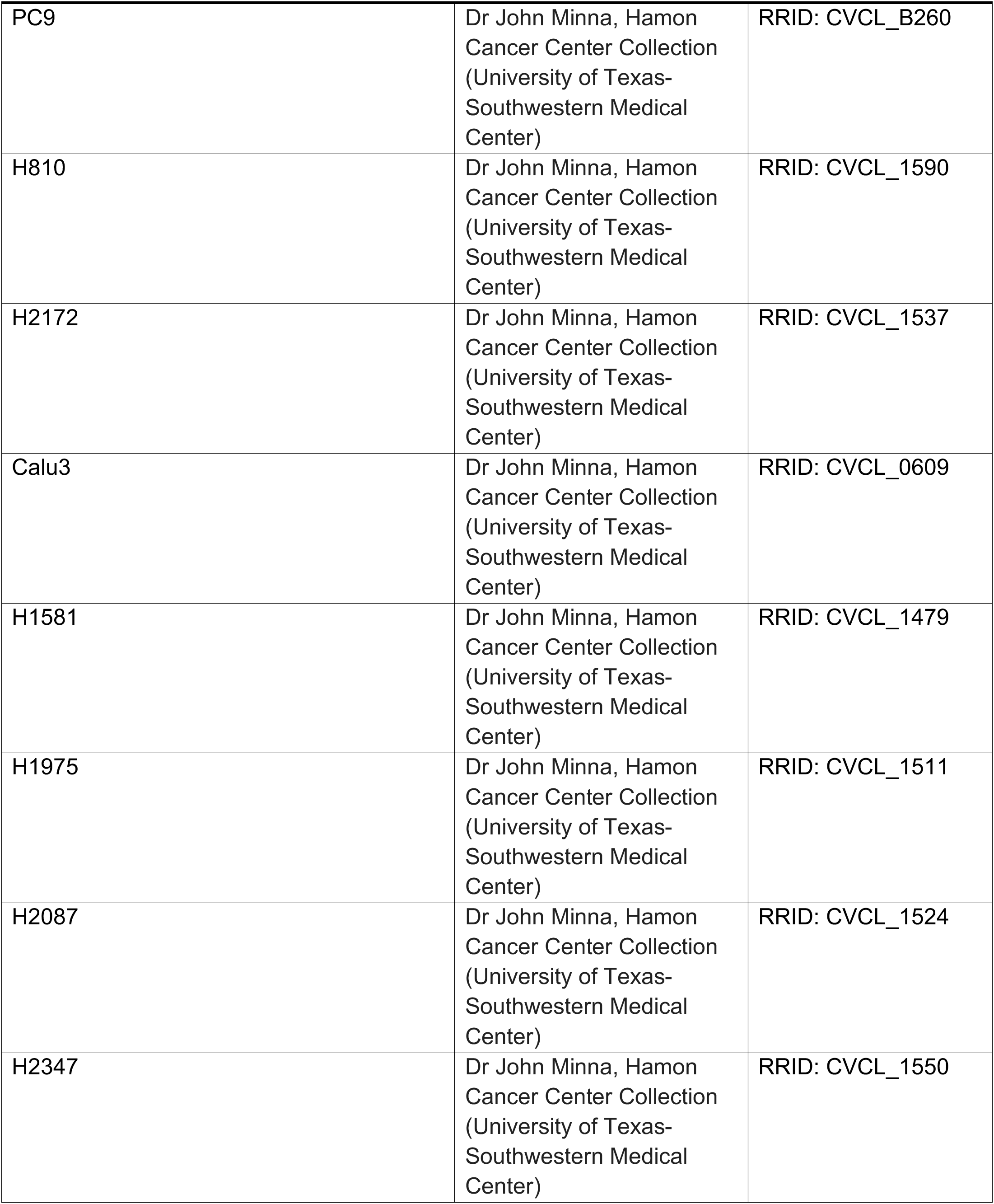
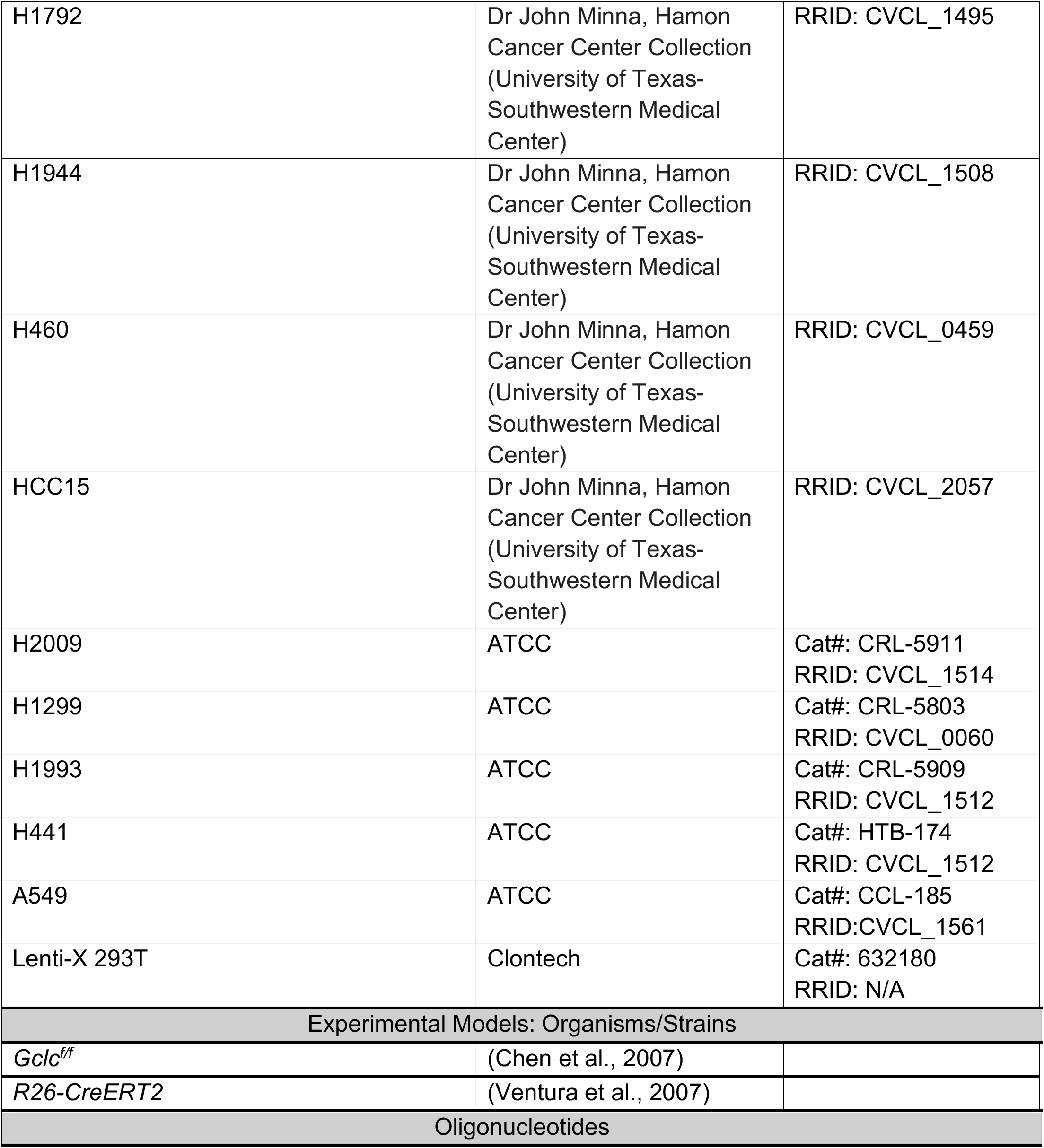
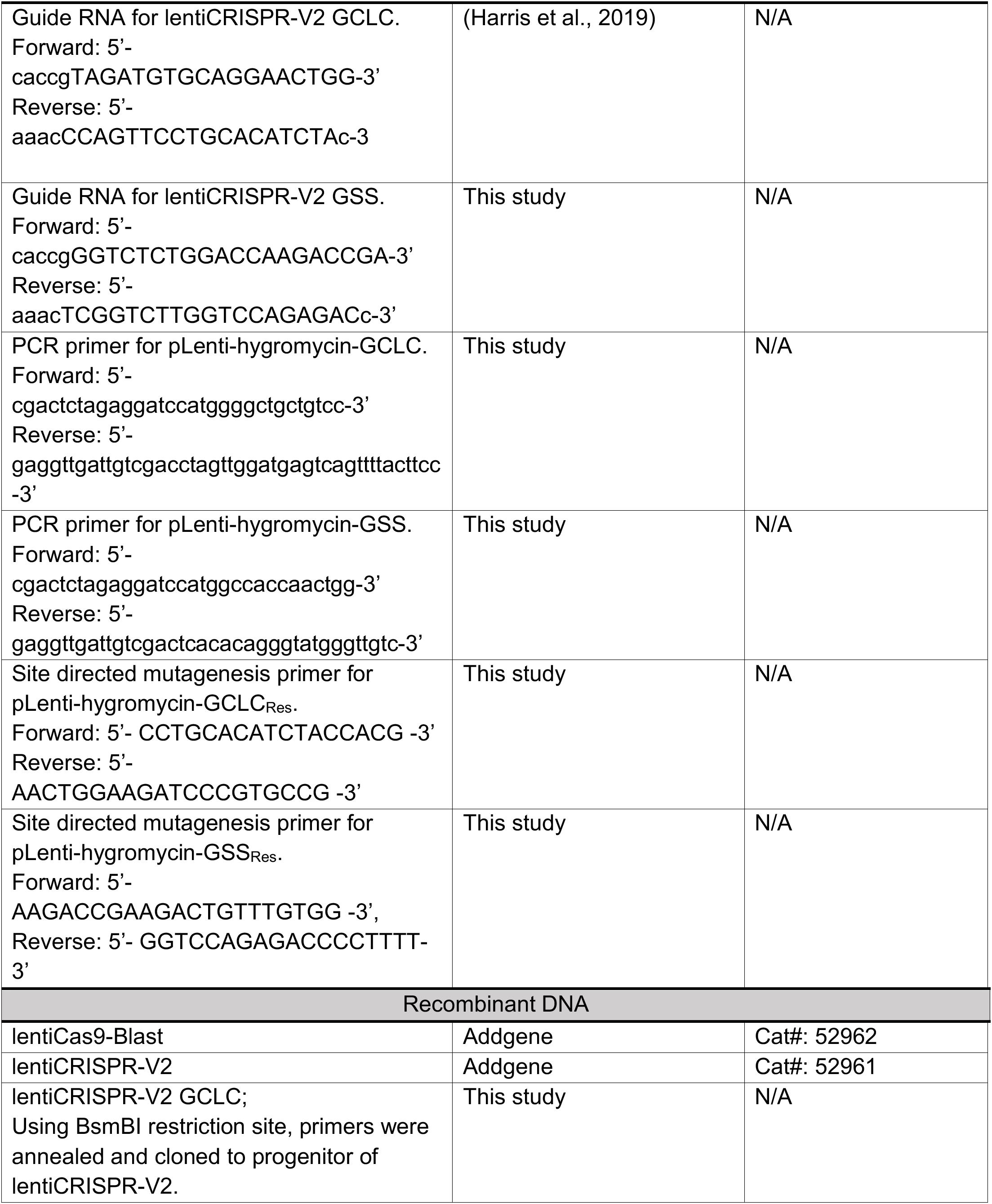
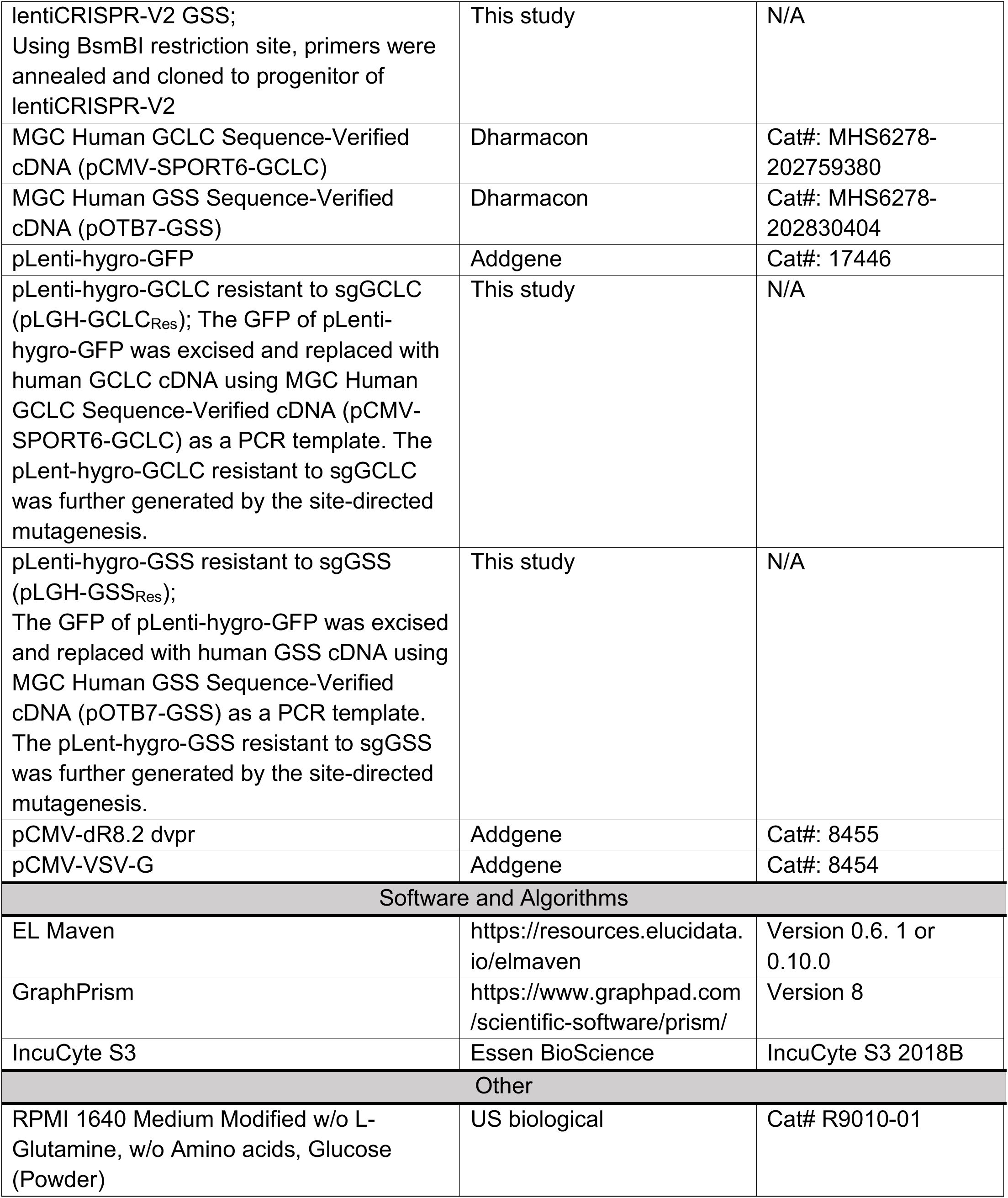
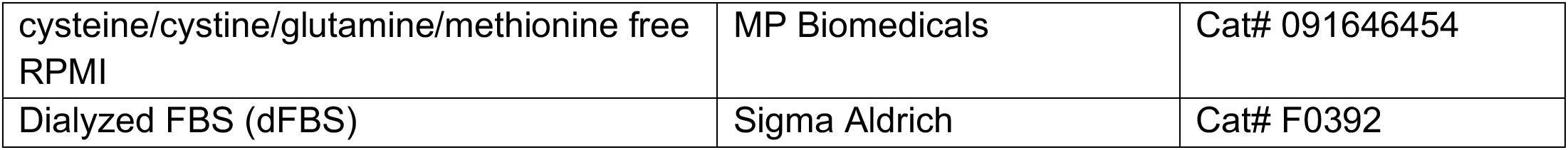

## Methods

### Animal Experiments

All animal studies were performed according to protocols approved by the Institutional Animal Care and Use Committee, the Standing Committee on Animals at Harvard University. *Gclc^f/f^* (Chen et al., 2007) and *R26-CreERT2* (Ventura et al., 2007) mice were bred to generate *Gclc^f/f^*; *R26-CreERT2* mice (C57B6 background). *Gclc^f/f^* (Gclc WT) and *Gclc^f/f^*; *R26-CreERT2* (Gclc KO) adult mice (aged 11-17 weeks old) were treated with tamoxifen (20 mg/ml; dissolved in corn oil) via daily intraperitoneal injection for 5 consecutive days at 160 mg/kg (tamoxifen/mouse body mass). Mice were humanely euthanized (isoflurane inhalation followed by cervical dislocation) 14-16 days later and necropsies were performed. Tissues (liver, kidney) were removed and serum was isolated; both were snap frozen on dry ice and stored at −80°C prior to analysis.

### Lentivirus generation

Lentiviruses were generated by overnight PEI transfection of 90% confluent Lenti-X 293T cells (Clonetech) with target lentiviral plasmid, and packaging plasmids pCMV-dR8.2 dvpr and pCMV-VSV-G in DMEM (10% FBS). The next day, the medium was changed to fresh DMEM (10% FBS). After 24 hrs, the first batch of virus contained medium was collected and filtered by 0.45 μm PES filter. The second batch of virus contained medium was collected as following above, further combined to the first batch and stored at −80°C until virus infection.

### Lentiviral infection of NSCLC cells

To increase of gene deletion efficiency in a polyclonal population, H1299 cells with stable *Streptococcus pyogenes* Cas9 expression (H1299^Cas9^) were established by lentiviral transduction and blasticidin selection (3 μg/mL) for 5 days, using lentiCas9-Blast (Greenfeld et al., 2015). To generate GCLC or GSS KO cells, H1299^Cas9^ and A549 cells were infected with the empty pLenti-CRISPR-V2 or pLenti-CRISPR-V2 encoding sgGCLC or sgGSS with 2 μg/mL of polybrene, followed by puromycin selection (1 μg/mL) for 4 days. To select single clones for GCLC or GSS KO, A549 cells were further diluted to 0.5 cells/well in 96 well dishes in RPMI supplemented with 1 μM of Fer-1. Cells were grown for 2 weeks, followed by expansion of clones. GCLC or GSS KO was verified by loss of GCLC or GSS protein via immunoblotting. Subsequently, cells were reconstituted with sgRNA-resistant cDNAs by infection with pLenti-hygro-GFP, pLenti-hygro-GCLC resistant to sgGCLC (pLGH-GCLC_Res_), or pLenti-hygro-GSS resistant to sgGSS (pLGH-GSS_Res_) followed by hygromycin selection (300 μg/mL) for 5 days.

### Dead cell measurement with Incucyte

Cells were plated in black walled 96 well pates at a density of 10,000 cells/well in 100 μL final volume. The next day, the medium was changed to 100 μL of experimental medium containing 20 nM of Sytox Green as follows: For the extracellular cystine starvation, cysteine/cystine, methionine and glutamine free RPMI (MP Biomedicals) was supplemented with 100 μM methionine and 10% dialyzed FBS (dFBS) containing 2 mM glutamine and/or 200 μM cystine as indicated. In addition, 10 μM of Ferrostatin-1 (Fer-1), 100 μM of DFO, 5 mM of GlutEE, 0.5 or 5 mM of AOA, or 100 μM of BSO was further supplemented as indicated. The number of dead cells and cell confluence were measured by IncuCyte S3 live-cell analysis system (Essen BioScience, Ann Arbor, MI, USA) in a humidified tissue culture incubator at 37°C with 5% CO_2_. Data were acquired by 10X objective lens in phase contrast and green fluorescence (excited wavelength: 460 nm, emission wavelength: 524 nm, acquisition time: 400 ms) channel. Images were acquired from each well at 3 hr intervals. Image and data processing were performed with IncuCyte S3 software (Essen BioScience, Ann Arbor, MI, USA). Dead cell number was normalized to cell confluence [Number of Sytox Green positive cells/mm^2^/cell confluent (%) of total image].

### Crystal Violet based cell viability assay

Cells were plated in 96-well plates at a density of 10,000-20,000 cells/well in 100 μL final volume. The next day, the medium was changed to 100 μL medium with different experimental conditions as indicated. At the indicated time points, cells were fixed with 4% Paraformaldehyde, stained with crystal violet, washed and dried. The crystal violet was dissolved in 10% acetic acid and the absorbance was measured by 600 nm wavelength. The relative cell number was normalized to control cells of each experimental set.

### Sample preparation for the liver tissue targeted metabolomics

The frozen liver tissue samples were pulverized using a pre-chilled BioPulverizer (59012MS, BioSpec), weighed frozen, and then placed on dry ice. The tissue metabolites were extracted in 80% MeOH at a final tissue concentration of 50mg/mL for 24 hrs at −80°C. After centrifugation (17000 g, 20 min, 4°C), the supernatant was analyzed by LC-HRMS.

### Sample preparation for the quantitative intracellular ^13^C_3_-serine tracing and cysteine quantification

NSCLC cells were plated in 6 well dishes and pre-conditioned in RPMI medium containing dFBS (10%) overnight. For serine tracing, RPMI 1640 Medium without glucose and amino acids (MP Biomedicals) was prepared from powder according the manufacturer’s instructions and supplemented with glucose and amino acids to meet the RPMI formulation with the exception of serine and cystine. The following day, the cells were quickly washed with 1 mL of warm serine/cystine-free RPMI medium, followed by feeding with ^13^C_3_-serine containing medium (serine/cystine-free RPMI + 10% dFBS + 1% Pen/Strep + 300 μM ^13^C_3_-Serine) lacking or supplemented with 200 μM cystine. After 4 hrs, the medium was aspirated and the cells were quickly washed with ice cold PBS. As described in previous study (Kang et al., 2019), the cellular metabolites were extracted and derivatized with 0.5 mL of ice-cold extraction solvent (80% MeOH:20% H2O containing 25 mM NEM and 10 mM ammonium formate, pH 7.0). The concentration of internal standards in the extraction solvent were as follows: 4.18 μM [^2^H_5_]-GSH-NEM, 2.49 μM [^13^C_3_, ^15^N]-serine and 2.48 uM [^13^C_2_, ^15^N]-glycine from METABOLOMICS AMINO ACID MIX STANDARD (Cambridge Isotope Laboratories). For cysteine quantification, the extraction solvent contained 10 μM [^13^C_2_, ^15^N]-cysteine-NEM. [^2^H_5_]-GSH-NEM and [^13^C_2_, ^15^N]-cysteine-NEM were pre-prepared by reaction of 50 mM NEM (10 mM ammonium formate, pH 7.0) for 30 min as previously described (Kang et al., 2019). After incubation on ice for 30 min, the NEM-derivatized metabolite extracts were cleared by centrifugation and the supernatant was analyzed by LC-HRMS at the positive mode. Cell volume and number were determined using a Scepter 2.0 cell counter (Millipore) and used to calculate the intracellular metabolite concentrations.

### Sample preparation for intracellular 2, 3, 3-^2^H_3_-serine tracing

A549 cells were prepared as described for ^13^C_3_-serine tracing but medium containing 300 μM [2, 3, 3-^2^H_3_-serine]-was used. Further, 0.5 μM of SHIN-1 (Ducker et al., 2017) was included in the cystine starved condition. After 12 hrs, the medium was aspirated, and cells were quickly washed with ice cold PBS, and cellular metabolites were extracted with 1 mL of 80% MeOH (−80°C, 15 min). After scraping, the metabolite extract was transferred into an Eppendorf tube and cleared by centrifugation (17000 g, 20 min, 4°C), followed by LC-HRMS analysis at the negative mode.

### Sample preparation for the intracellular non-targeted metabolomics, and Glycine and Glutamate quantification

NSCLC cells were plated in 6 well dishes so they were 70% confluent at extraction and preconditioned in RPMI medium containing dFBS (10%) overnight. The following day, the medium was aspirated and the cells were quickly washed with 1 mL of RPMI (10% dFBS, 1% P/S), followed by feeding with conditioning medium as indicated. 1 mL of the medium supernatant was collected to assay glutamate as indicated. For the non-targeted metabolomics approach, medium was aspirated and cells were quickly washed with ice cold PBS, followed by extraction of cellular metabolites with 0.5 mL of 80% MeOH (−80°C, 15 min). For intracellular glutamate and glycine quantification, the extraction solvent also contained 2.49 uM [^13^C_5_, ^15^N]-glutamate and 2.48 uM [^13^C_2_, 15N]-glycine from METABOLOMICS AMINO ACID MIX STANDARD (Cambridge Isotope Laboratories). After scraping, the metabolite extract was transferred into an Eppendorf tube and cleared by centrifugation (17000 g, 20 min, 4°C), followed by LC-HRMS analysis in the negative or positive mode. For glutamate and glycine quantification, cell volume and number were further determined using a Scepter 2.0 cell counter (Millipore).

### Quantitation of extracellular glutamate exportation

The extracellular medium collected above was transferred to the 96 well plates and the medium glutamate levels was measured by YSI 2900 (Yellow springs, OH, USA) using 2755 glutamate standard (5 mM) The extracellular glutamate secretion rate (nmol/μL of cell volume/hrs) determined from cell volume measurements above.

### LC-MS analysis

The LC-MS condition was identical with previously established method (Kang et al., 2019). For the chromatographic metabolite separation, the Vanquish UPLC systems coupled to a Q Exactive HF (QE-HF) mass spectrometer equipped with HESI (Thermo Fisher Scientific, Waltham, MA). The column was a SeQuant ZIC-pHILIC LC column, 5 μm, 150 × 4.6 mm (MilliporeSigma, Burlington, MA) with a SeQuant ZIC-pHILIC guard column, 20 × 4.6 mm (MilliporeSigma, Burlington, MA). Mobile phase A was 10 mM (NH_4_)_2_CO_3_ and 0.05% NH_4_OH in H_2_O while mobile phase B was 100% ACN. The column chamber temperature was set to 30°C. The mobile phases were eluted as following gradient condition. 0-13min: 80% to 20% of mobile phase B, 13-15min: 20% of mobile phase B. The ESI ionization mode was positive or negative. The MS scan range (m/z) was set to 60-900. The mass resolution was 120,000 and the AGC target was 3 × 10^6^. The capillary voltage and capillary temperature were set to 3.5 KV and 320°C, respectively. The 5 μL of sample was loaded. For targeted metabolomics, the LC-MS peaks were manually identified and integrated by EL-Maven (Version 0. 6. 1) by matching with a previously established in-house library (Kang et al., 2019). The peak area of target metabolites was further normalized by the median value of identified metabolite peak areas or the peak area of stable isotope labeled internal standards for further quantification as previously described (Bennett et al., 2008). For the non-targeted metabolomics approach, the LC-MS peaks were automatically extracted and aligned using the Automated Feature Detection function of ELMaven. After the normalization with median value of the intensities of LC-MS peaks, the statistical analysis was conducted. The γ-glutamyl peptides peaks were putatively identified by matching m/z value with online HMDB database (http://www.hmdb.ca), and further confirmed by matching with m/z value and retention time of authentic standards. The standards of γ-glutamyl threonine and γ-glutamyl-alanyl-glycine were not available and were instead validated by stable isotope labeled metabolite tracing (Figure S4A and B), as described in result.

### ROS measurement by CellRox green

NSCLC cells were plated at 70,000 cells/well to 24 well dishes and pre-conditioned in RPMI medium containing dFBS (10%) overnight. The following day, the medium was aspirated and the cells were quickly washed with the warm PBS followed by feeding with the indicated medium conditions. The CellRox green was added to the cells at a final concentration of 5 μM for the final 30 min of the experiment. Cells were washed with PBS, detached with trypsin, and transferred to Eppendorf tubes. After spin down (10sec, 17,000g), the supernatant was aspirated and the pellet was re-suspended in 500 μL of PBS and filtered into FACS tubes. The mean fluorescence intensity (MFI) of Green fluorescence from the 4,000 discrete events was determined by Accuri™ C6 Flow cytometry (BD biosciences, An Arbor, Mi, USA). The MFI value was obtained by the Accuri™ C6 software (BD biosciences, An Arbor, Mi, USA) and further normalized by the control of experimental set.

### Immunoblotting

Cell lysates were prepared in RIPA buffer (20 mM Tris-HCl, pH7.5; 150 mM NaCl, 1 mM EDTA, 1 mM EGTA, 1% NP-40, 1% sodium deoxycholate) containing protease inhibitors. After protein quantification using Biorad DC assay, the samples were mixed with reducing buffer (v/v, 5:1) containing 2-mercaptoethanol. The proteins were separated by SDS-PAGE using NuPAGE [4-12% Bis-Tris gels (Invitrogen)] and transferred to 0.45 um Nitrocellulose membrane (GE Healthcare). The membrane was blocked by 5% non-fat milk in TBST for 15 min, and the primary antibodies were incubated overnight in blocking buffer as follows: GCLC - 1/1000 dilution with 5% non-fat milk in TBST; GSS - 1/2000 dilution with 5% non-fat milk in TBST; HSP90 - 1/5000 dilution with 5% non-fat milk in TBST. After wash membrane 3 times using TBST for 10 min of each, the 10,000 times diluted secondary antibody in 5% non-fat milk in TBST (goat anti-rabbit or goat anti-mouse, Jackson ImmunoResearch) were attached to membrane for 1 hr. After wash the membrane in TBST for 3 times for 10 min of each, the enhanced chemo-luminescence signal was measured by exposing to the x-ray film followed by fixing and developing.

### Statistical analysis

Statistical analyses were conducted with Graph Pad Prism 8. For the comparison of two groups, two-tailed Student’s t-test was used. For the comparison of more than 3 experimental groups, one-way ANOVA was used with Bonferroni’s multiple comparison test.

## Lead Contact

Further information and requests for resources and reagents should be directed to and will be fulfilled by the Lead Contact, Dr. Gina M. DeNicola

## Acknowledgement

We would like to thank Dr. Joshua D. Rabinowitz for SHIN-1, Dr. Vince Luca, Dr. David Gonzalez-Perez and Elliot Medina for flow cytometry assistance, Dr. Min Liu for LC-MS assistance, Chen Tingan for Incucyte assistance, and all members of the DeNicola laboratory for their very helpful discussions. This work was supported by grants from the NIH/NCI (R37-CA230042) to G.M.D, the Ludwig Center at Harvard to I.S.H., the AACR-Takeda Oncology Lung Cancer Research Fellowship (19-40-38-KANG) to Y.P.K., a National Pancreas Foundation grant to C.J. This work was also supported by the Analytic Microscopy and the Proteomics/Metabolomics Cores, which are funded in part by Moffitt’s Cancer Center Support Grant (NCI, P30-CA076292), and grants from the Moffitt Foundation, and a Florida Bankhead-Coley grant (06BS-02–9614) to the Proteomics/Metabolomics core.

## Author Contributions

Y.P.K and G.M.D. designed the study and interpreted experimental results. Y.P.K performed all metabolomics experiments, Y.P.K performed western blotting and cell viability experiments with assistance from A.M-M., Y.P.K. generated KO cell lines with assistance from C.J., I.S.H. generated Gclc knockout mice and collected tissues, Y.P.K and G.M.D wrote the manuscript and all authors commented on it.

## Declaration of Interests

The authors declare no competing financial interests. I.S.H. is a consultant for Ono Pharma USA, who had no role in funding or design of this study.

## Supplementary Figure legends

**Figure S1.**
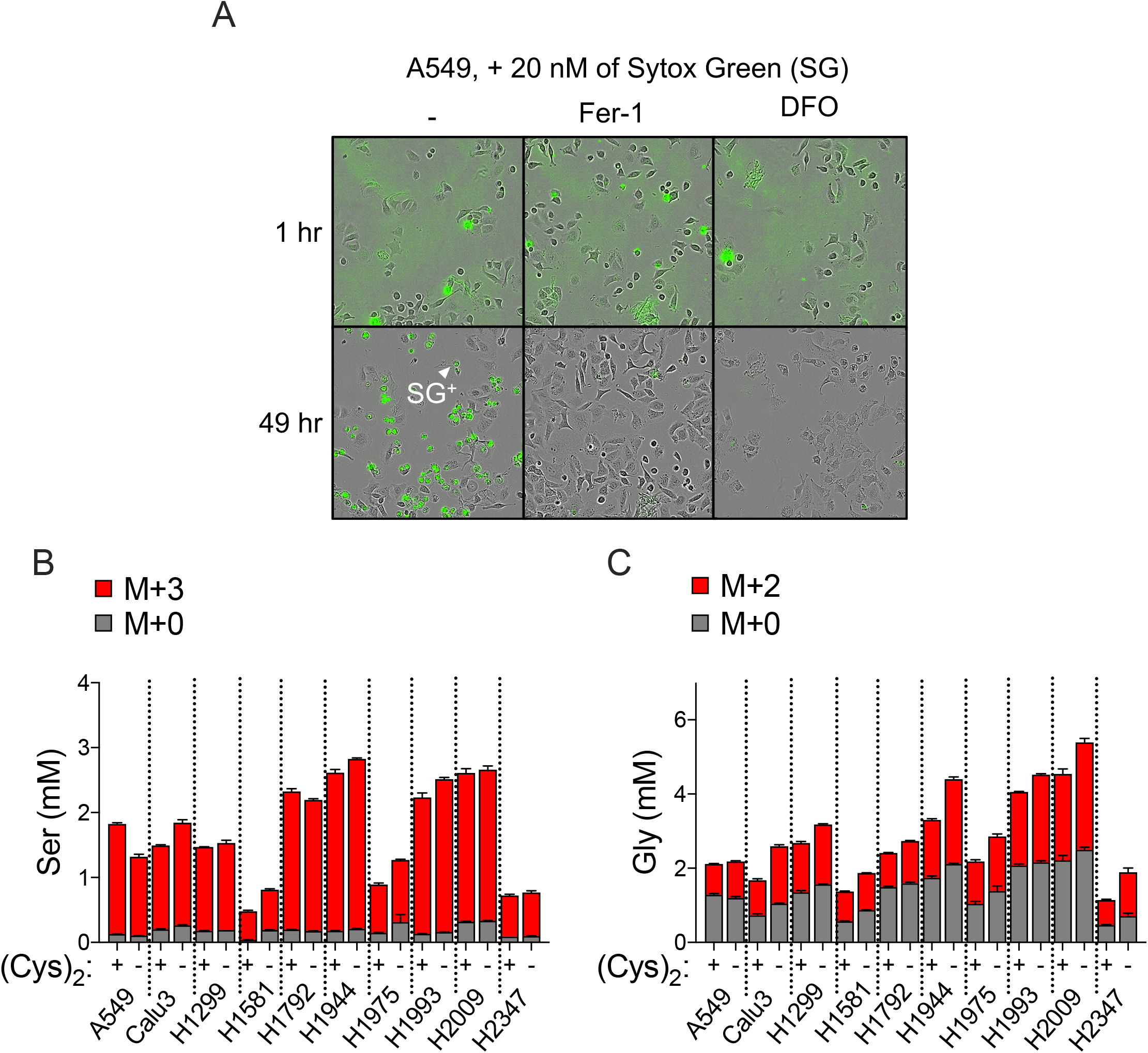
Cysteine starvation induces ferroptosis and impairs GSH synthesis. (Related to Figure 1) (A) Representative image of sytox green (SG) stained A549 cells under cystine starvation conditions for 1 or 49 hours treated with Vehicle (0.1% DMSO), Fer-1 (10 μM), or DFO (100 μM). Images show the same well position at the two different time points (1 or 49 hrs). A representative sytox green positive (SG^+^) cell is indicated with a white arrow. (B) Quantitative [^13^C_3_]-Serine labeling of (B) serine (Ser) and (C) glycine (Gly) following media change to cystine starved (-) or replete (+) conditions for 4 hrs. (N=3 biological replicates). Data shown as mean ± SEM.

**Figure S2.**
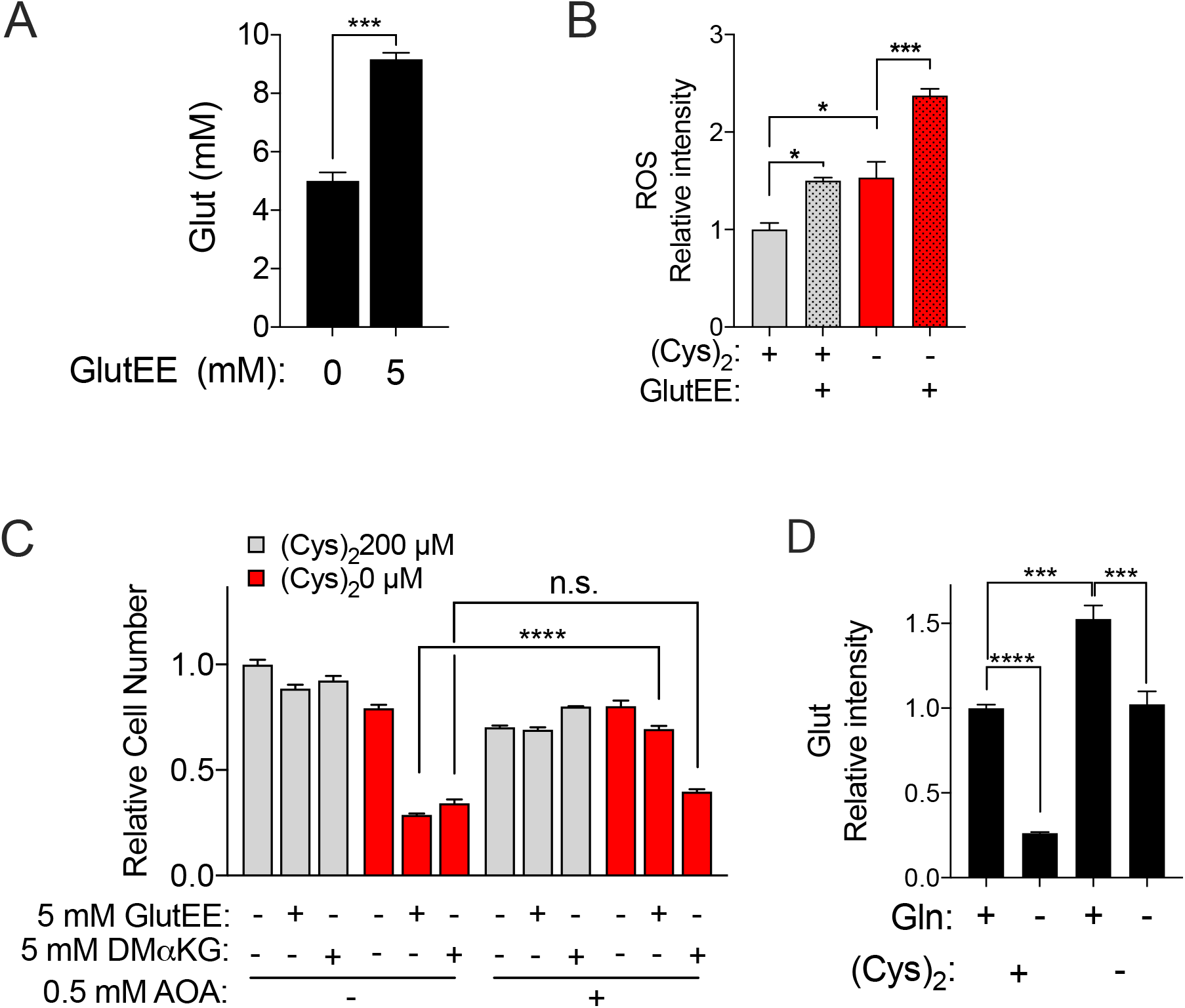
Inhibition of GSH synthesis promotes ferroptosis via glutamate accumulation. (Related to Figure 2) (A) Quantitative measurement of intracellular glutamate levels in A549 cells cultured with the indicated concentration of GlutEE (N=3). (B) Measurement of intracellular ROS levels under cystine starved or replete condition treated with vehicle or GlutEE (5 mM) for 4 hrs (N=3). Data was normalized to the mean value of cells under cystine replete conditions without GlutEE. (C) Measurement of cell number under cystine starved or replete condition with vehicle, GlutEE (5 mM), dimethyl alpha-ketoglutarate (DMαKG, 5 mM), and/or AOA (0.5 mM) as indicated for 17 hrs (N=3). Data was normalized to the mean value of vehicle-treated cells under cystine replete conditions. (D) Measurement of intracellular glutamate levels under cystine starved or replete conditions cultured with or without media glutamine for 3 hrs (N=3). For A-D, data are shown as mean ± SEM. N is number of biological replicates. n.s., not significant; *P<0.05, ***P<0.001, and ****P<0.0001. For A and C, an unpaired two-tailed t test was used for statistical analyses. For B and D, a one-way ANOVA with Bonferroni’s multiple comparison test was used.

**Figure S3.**
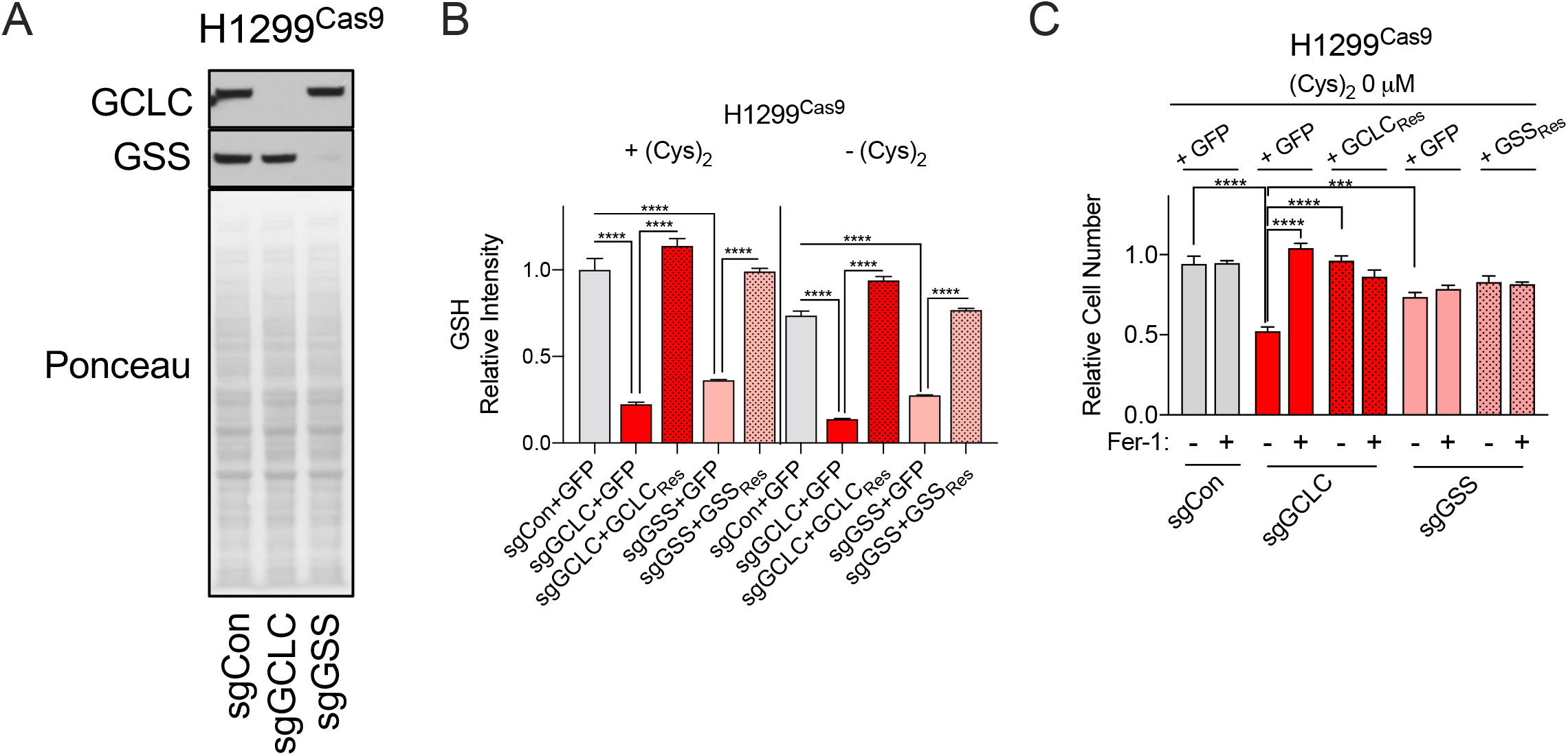
GCLC prevents ferroptosis independent of GSH production. (Related to Figure 3). (A) Representative immunoblots of GCLC and GSS from H1299^Cas9^ cells transfected with LentiCrisprV2 with control sgRNA (sgCon) or sgRNA targeting GCLC (sgGCLC) and GSS (sgGSS). Ponceau staining was used for the loading control. (B) Intracellular GSH levels of H1299^Cas9^ infected with sgCon, sgGCLC or sgGSS reconstituted with GFP (+GFP), sgRNA-resistant GCLC (+GCLC_Res_), or sgRNA-resistant GSS (+GSS_Res_) as indicated under cystine replete or starved condition for 3 hrs (N=3). Data was normalized to the mean value of sgCon with GFP cells under cystine replete conditions. (C) Relative cell number of H1299^Cas9^ infected with sgCon, sgGCLC or sgGSS reconstituted with GFP, GCLC_Res_, or GSS_Res_ as indicated under cystine starved condition treated with vehicle (0.1% DMSO), Fer-1 (10 μM), DFO (100 μM) for 13 hrs (N=3). Data was normalized to the mean value of vehicle-treated cystine replete conditions. Data shown as mean ± SEM (B and C). N is number of biological replicates. ***P<0.001, ****P<0.0001. One-way ANOVA with Bonferroni’s multiple comparison test was used to determined statistical significance for B and C.

**Figure S4.**
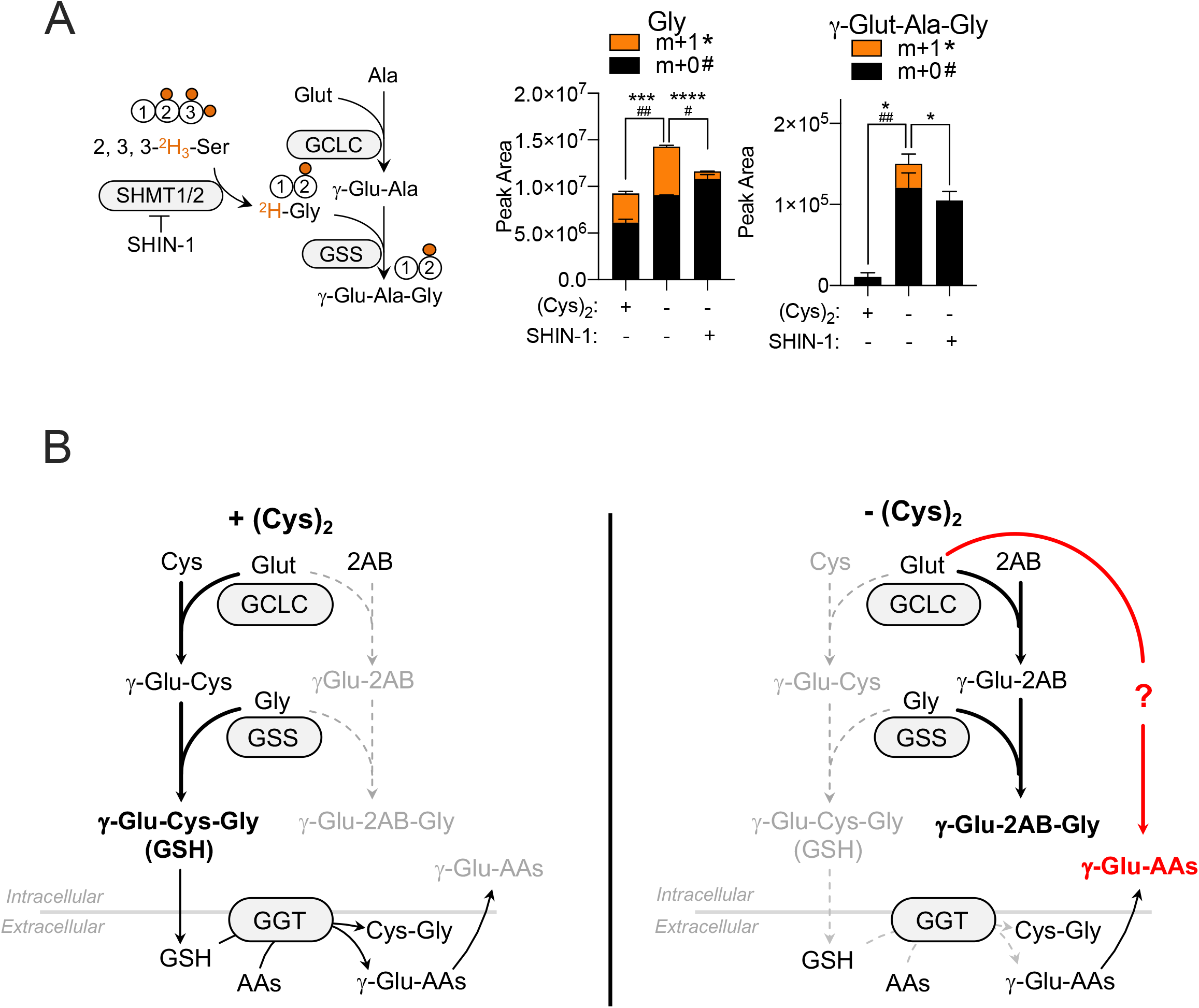
Cystine starvation promotes γ-Glut-dipeptide production by GCLC. (Related to Figure 4). (A) 2, 3, 3-^2^H_3_-Ser tracing of A549 cells under cystine replete or starved condition in the presence and absence of SHIN-1 (0.5 uM) as indicated for 12 hrs (N=3). (B) Schematic depiction of the mechanism of γ-glutamyl dipeptides synthesis under cystine replete and starved conditions. While γ-glutamyl transferase (GGT) can generate of γ-glutamyl dipeptides from glutathione, GCLC can generate γ-glutamyl dipeptides from glutamate and amino acids under cystine starved conditions.

**Figure 5S.**
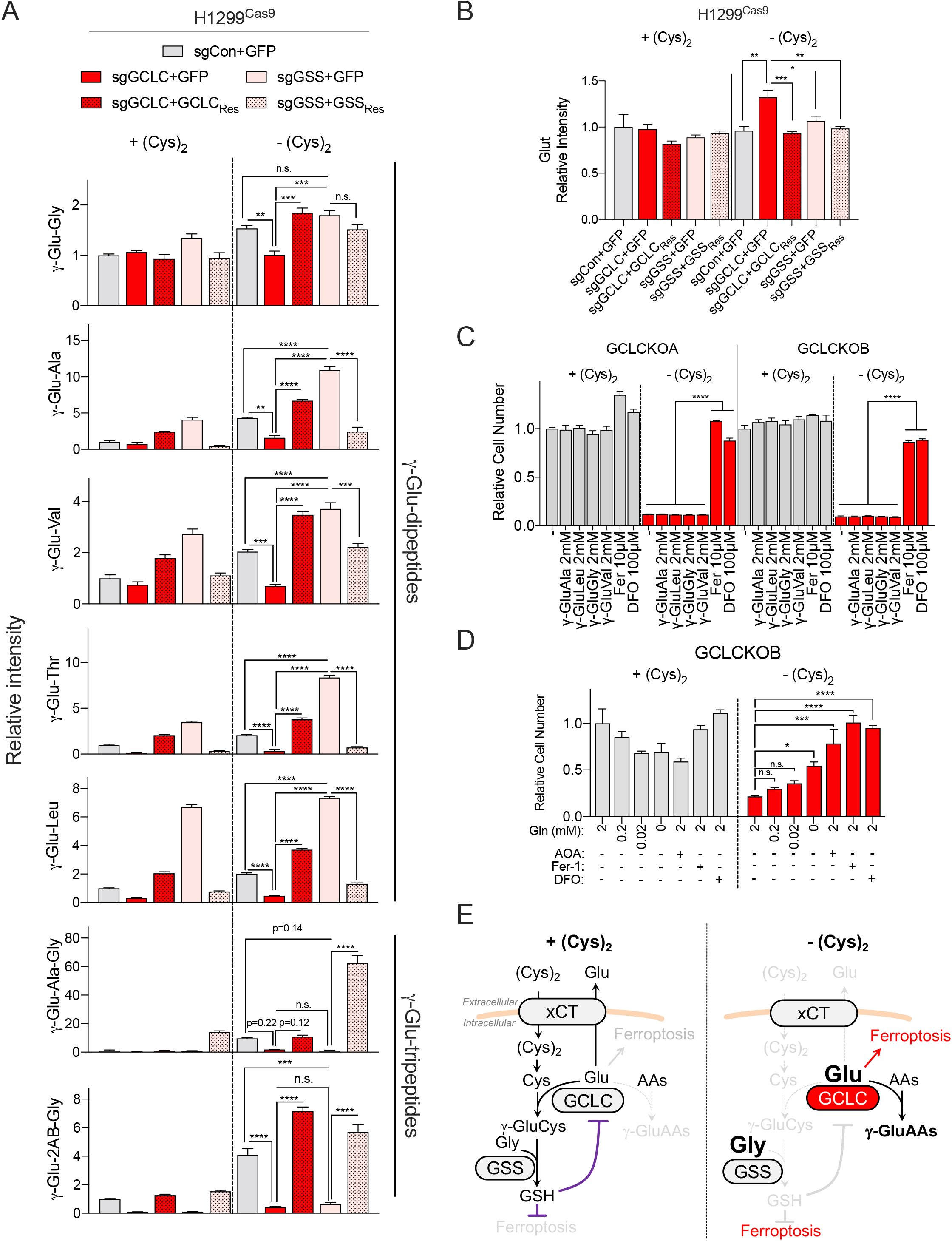
γ-glutamyl peptide synthesis by GCLC scavenges glutamate to protect against ferroptosis (Related to Figure 5). (A-B) Measurement of intracellular (A) γ-glutamyl-peptide and (B) Glut levels of H1299^Cas9^ infected with sgCon, sgGCLC or sgGSS reconstituted with GFP (+GFP), sgRNA-resistant GCLC (+GCLC_Res_), or sgRNA-resistant GSS (+GSS_Res_) as indicated under cystine replete or starved conditions for 3 hrs (N=3). Data was normalized to the mean value of sgCon +GFP cells under cystine replete conditions. (C) Analysis of cell number of GCLC KO A549 clones following cystine replete or starved conditions and treatment with vehicle, γ-glutamyl dipeptides (γ-Glu-Ala, γ-Glu-Leu, γ-Glu-Gly, and γ-Glu-Val; 2 mM), Fer-1 (10 uM), or DFO (100 uM) as indicated for 14 hrs (N=3). The data were normalized to the mean value of cystine replete, vehicle treated conditions for each clone. (D) Analysis of cell number of GCLC KO A549 cells following cystine replete or starved conditions and treatment with vehicle, glutamine starvation (0, 0.02, 0.2 mM), AOA (5 mM), Fer-1 (10 uM), or DFO (100 uM) as indicated for 12 hrs (N=3). The data were normalized to the mean value of cystine replete, vehicle treated conditions for each clone. For A-D, data are shown as mean ± SEM. N is number of biological replicates. n.s., not significant; *P<0.05, ***P<0.001, ****P<0.0001. For AD, one-way ANOVA with Bonferroni’s multiple comparison test was used for statistical analyses. (E) Schematic depiction of GSH-independent GCLC function that prevents ferroptosis under cystine starvation via depletion of glutamate.

## References

1. Anderson, M.E. (1998). Glutathione: an overview of biosynthesis and modulation. Chemico-biological interactions 111, 1–14.

2. Balendiran, G.K., Dabur, R., and Fraser, D. (2004). The role of glutathione in cancer. Cell Biochemistry and Function: Cellular biochemistry and its modulation by active agents or disease 22, 343–352.

3. Beatty, P.W., and Reed, D.J. (1980). Involvement of the cystathionine pathway in the biosynthesis of glutathione by isolated rat hepatocytes. Archives of biochemistry and biophysics 204, 80–87.

4. Bennett, B.D., Yuan, J., Kimball, E.H., and Rabinowitz, J.D. (2008). Absolute quantitation of intracellular metabolite concentrations by an isotope ratio-based approach. Nature protocols 3, 1299.

5. Cao, J.Y., and Dixon, S.J. (2016). Mechanisms of ferroptosis. Cellular and Molecular Life Sciences 73, 2195–2209.

6. Chen, Y., Yang, Y., Miller, M.L., Shen, D., Shertzer, H.G., Stringer, K.F., Wang, B., Schneider, S.N., Nebert, D.W., and Dalton, T.P. (2007). Hepatocyte-specific Gclc deletion leads to rapid onset of steatosis with mitochondrial injury and liver failure. Hepatology 45, 1118–1128.

7. Conrad, M., and Friedmann Angeli, J.P. (2015). Glutathione peroxidase 4 (Gpx4) and ferroptosis: what’s so special about it? Molecular & cellular oncology 2, e995047.

8. Cramer, S.L., Saha, A., Liu, J., Tadi, S., Tiziani, S., Yan, W., Triplett, K., Lamb, C., Alters, S.E., and Rowlinson, S. (2017). Systemic depletion of L-cyst (e) ine with cyst (e) inase increases reactive oxygen species and suppresses tumor growth. Nature medicine 23, 120.

9. Dixon, S.J., Lemberg, K.M., Lamprecht, M.R., Skouta, R., Zaitsev, E.M., Gleason, C.E., Patel, D.N., Bauer, A.J., Cantley, A.M., and Yang, W.S. (2012). Ferroptosis: an iron-dependent form of nonapoptotic cell death. Cell 149, 1060–1072.

10. Dixon, S.J., and Stockwell, B.R. (2019). The hallmarks of ferroptosis. Annual Review of Cancer Biology 3, 35–54.

11. Dixon, S.J., Winter, G.E., Musavi, L.S., Lee, E.D., Snijder, B., Rebsamen, M., Superti-Furga, G., and Stockwell, B.R. (2015). Human haploid cell genetics reveals roles for lipid metabolism genes in nonapoptotic cell death. ACS chemical biology 10, 1604–1609.

12. Doll, S., Proneth, B., Tyurina, Y.Y., Panzilius, E., Kobayashi, S., Ingold, I., Irmler, M., Beckers, J., Aichler, M., and Walch, A. (2017). ACSL4 dictates ferroptosis sensitivity by shaping cellular lipid composition. Nature chemical biology 13, 91.

13. Ducker, G.S., Ghergurovich, J.M., Mainolfi, N., Suri, V., Jeong, S.K., Hsin-Jung Li, S., Friedman, A., Manfredi, M.G., Gitai, Z., Kim, H., et al. (2017). Human SHMT inhibitors reveal defective glycine import as a targetable metabolic vulnerability of diffuse large B-cell lymphoma. Proceedings of the National Academy of Sciences 114, 11404–11409.

14. Furuyama, K., and Sassa, S. (2000). Interaction between succinyl CoA synthetase and the heme-biosynthetic enzyme ALAS-E is disrupted in sideroblastic anemia. J Clin Invest 105, 757–764.

15. Gao, M., Monian, P., Quadri, N., Ramasamy, R., and Jiang, X. (2015). Glutaminolysis and transferrin regulate ferroptosis. Molecular cell 59, 298–308.

16. Gao, M., Yi, J., Zhu, J., Minikes, A.M., Monian, P., Thompson, C.B., and Jiang, X. (2019). Role of mitochondria in ferroptosis. Molecular cell 73, 354–363. e353.

17. Greenfeld, H., Takasaki, K., Walsh, M.J., Ersing, I., Bernhardt, K., Ma, Y., Fu, B., Ashbaugh, C.W., Cabo, J., and Mollo, S.B. (2015). TRAF1 coordinates polyubiquitin signaling to enhance Epstein-Barr virus LMP1-mediated growth and survival pathway activation. PLoS pathogens 11.

18. Habib, E., Linher-Melville, K., Lin, H.-X., and Singh, G. (2015). Expression of xCT and activity of system xc-are regulated by NRF2 in human breast cancer cells in response to oxidative stress. Redox biology 5, 33–42.

19. Hanigan, M.H., and Pitot, H.C. (1985). Gamma-glutamyl transpeptidase–its role in hepatocarcinogenesis. Carcinogenesis 6, 165–172.

20. Harris, I.S., and DeNicola, G.M. (2020). The Complex Interplay between Antioxidants and ROS in Cancer. Trends in Cell Biology.

21. Harris, I.S., Endress, J.E., Coloff, J.L., Selfors, L.M., McBrayer, S.K., Rosenbluth, J.M., Takahashi, N., Dhakal, S., Koduri, V., and Oser, M.G. (2019). Deubiquitinases maintain protein homeostasis and survival of cancer cells upon glutathione depletion. Cell metabolism 29, 1166–1181. e1166.

22. Harris, I.S., Treloar, A.E., Inoue, S., Sasaki, M., Gorrini, C., Lee, K.C., Yung, K.Y., Brenner, D., Knobbe-Thomsen, C.B., and Cox, M.A. (2015). Glutathione and thioredoxin antioxidant pathways synergize to drive cancer initiation and progression. Cancer cell 27, 211–222.

23. Huang, C.-S., Moore, W.R., and Meister, A. (1988). On the active site thiol of gamma-glutamylcysteine synthetase: relationships to catalysis, inhibition, and regulation. Proceedings of the National Academy of Sciences 85, 2464–2468.

24. Huang, Z.-Z., Chen, C., Zeng, Z., Yang, H., Oh, J., Chen, L., and Lu, S.C. (2001). Mechanism and significance of increased glutathione level in human hepatocellular carcinoma and liver regeneration. The FASEB Journal 15, 19–21.

25. Ji, X., Qian, J., Rahman, S.J., Siska, P.J., Zou, Y., Harris, B.K., Hoeksema, M.D., Trenary, I.A., Heidi, C., and Eisenberg, R. (2018). xCT (SLC7A11)-mediated metabolic reprogramming promotes non-small cell lung cancer progression. Oncogene 37, 5007–5019.

26. Jiang, L., Kon, N., Li, T., Wang, S.-J., Su, T., Hibshoosh, H., Baer, R., and Gu, W. (2015). Ferroptosis as a p53-mediated activity during tumour suppression. Nature 520, 57–62.

27. Kang, Y.P., Torrente, L., Falzone, A., Elkins, C.M., Liu, M., Asara, J.M., Dibble, C.C., and DeNicola, G.M. (2019). Cysteine dioxygenase 1 is a metabolic liability for non-small cell lung cancer. Elife 8, e45572.

28. Kobayashi, S., Ikeda, Y., Shigeno, Y., Konno, H., and Fujii, J. (2020). γ-Glutamylcysteine synthetase and γ-glutamyl transferase as differential enzymatic sources of γ-glutamylpeptides in mice. Amino Acids, 1–12.

29. Lim, J.K., Delaidelli, A., Minaker, S.W., Zhang, H.-F., Colovic, M., Yang, H., Negri, G.L., von Karstedt, S., Lockwood, W.W., and Schaffer, P. (2019). Cystine/glutamate antiporter xCT (SLC7A11) facilitates oncogenic RAS transformation by preserving intracellular redox balance. Proceedings of the National Academy of Sciences 116, 9433–9442.

30. Mancias, J.D., Wang, X., Gygi, S.P., Harper, J.W., and Kimmelman, A.C. (2014). Quantitative proteomics identifies NCOA4 as the cargo receptor mediating ferritinophagy. Nature 509, 105–109.

31. Martinez-Reyes, I., Diebold, L.P., Kong, H., Schieber, M., Huang, H., Hensley, C.T., Mehta, M.M., Wang, T., Santos, J.H., Woychik, R., et al. (2016). TCA Cycle and Mitochondrial Membrane Potential Are Necessary for Diverse Biological Functions. Mol Cell 61, 199–209.

32. Ookhtens, M., and Kaplowitz, N. (1998). Role of the liver in interorgan homeostasis of glutathione and cyst (e) ine. In Seminars in liver disease (© 1998 by Thieme Medical Publishers, Inc.), pp. 313–329.

33. Oppenheimer, L., Wellner, V.P., Griffith, O.W., and Meister, A. (1979). Glutathione synthetase. Purification from rat kidney and mapping of the substrate binding sites. Journal of Biological Chemistry 254, 5184–5190.

34. Penno, A., Reilly, M.M., Houlden, H., Laurá, M., Rentsch, K., Niederkofler, V., Stoeckli, E.T., Nicholson, G., Eichler, F., and Brown, R.H. (2010). Hereditary sensory neuropathy type 1 is caused by the accumulation of two neurotoxic sphingolipids. Journal of biological chemistry 285, 11178–11187.

35. Rao, A.M., Drake, M.R., and Stipanuk, M.H. (1990). Role of the transsulfuration pathway and of γ-cystathionase activity in the formation of cysteine and sulfate from methionine in rat hepatocytes. The Journal of nutrition 120, 837–845.

36. Reed, D.J., and Orrenius, S. (1977). The role of methionine in glutathione biosynthesis by isolated hepatocytes. Biochemical and biophysical research communications 77, 1257–1264.

37. Richman, P., and Meister, A. (1975). Regulation of gamma-glutamyl-cysteine synthetase by nonallosteric feedback inhibition by glutathione. Journal of Biological Chemistry 250, 1422–1426.

38. Ristoff, E., and Larsson, A. (2007). Inborn errors in the metabolism of glutathione. Orphanet journal of rare diseases 2, 16.

39. Rouault, T.A. (2012). Biogenesis of iron-sulfur clusters in mammalian cells: new insights and relevance to human disease. Dis Model Mech 5, 155–164.

40. Sasaki, H., Sato, H., Kuriyama-Matsumura, K., Sato, K., Maebara, K., Wang, H., Tamba, M., Itoh, K., Yamamoto, M., and Bannai, S. (2002). Electrophile response element-mediated induction of the cystine/glutamate exchange transporter gene expression. Journal of Biological Chemistry 277, 44765–44771.

41. Shin, C.-S., Mishra, P., Watrous, J.D., Carelli, V., D’Aurelio, M., Jain, M., and Chan, D.C. (2017). The glutamate/cystine xCT antiporter antagonizes glutamine metabolism and reduces nutrient flexibility. Nature communications 8, 1–11.

42. Sofyanovich, O.A., Nishiuchi, H., Yamagishi, K., Matrosova, E.V., and Serebrianyi, V.A. (2019). Multiple pathways for the formation of the γ-glutamyl peptides γ-glutamyl-valine and γ-glutamyl-valyl-glycine in Saccharomyces cerevisiae. PloS one 14.

43. Soga, T., Sugimoto, M., Honma, M., Mori, M., Igarashi, K., Kashikura, K., Ikeda, S., Hirayama, A., Yamamoto, T., and Yoshida, H. (2011). Serum metabolomics reveals γ-glutamyl dipeptides as biomarkers for discrimination among different forms of liver disease. Journal of hepatology 55, 896–905.

44. Soini, Y., Näpänkangas, U., Järvinen, K., Kaarteenaho-Wiik, R., Pääkkö, P., and Kinnula, V.L. (2001). Expression of γ-glutamyl cysteine synthetase in nonsmall cell lung carcinoma. Cancer: Interdisciplinary International Journal of the American Cancer Society 92, 2911–2919.

45. Solmonson, A., and DeBerardinis, R.J. (2018). Lipoic acid metabolism and mitochondrial redox regulation. J Biol Chem 293, 7522–7530.

46. Stipanuk, M., Ueki, I., Dominy, J., Simmons, C., and Hirschberger, L. (2009). Cysteine dioxygenase: a robust system for regulation of cellular cysteine levels. Amino acids 37, 55.

47. Stipanuk, M.H., Dominy Jr, J.E., Lee, J.-I., and Coloso, R.M. (2006). Mammalian cysteine metabolism: new insights into regulation of cysteine metabolism. The Journal of nutrition 136, 1652S–1659S.

48. Sun, J., Zhou, C., Ma, Q., Chen, W., Atyah, M., Yin, Y., Fu, P., Liu, S., Hu, B., and Ren, N. (2019). High GCLC level in tumor tissues is associated with poor prognosis of hepatocellular carcinoma after curative resection. Journal of Cancer 10, 3333.

49. Takeuchi, S., Wada, K., Toyooka, T., Shinomiya, N., Shimazaki, H., Nakanishi, K., Nagatani, K., Otani, N., Osada, H., and Uozumi, Y. (2013). Increased xCT expression correlates with tumor invasion and outcome in patients with glioblastomas. Neurosurgery 72, 33–41.

50. Tatebe, S., Unate, H., Sinicrope, F.A., Sakatani, T., Sugamura, K., Makino, M., Ito, H., Savaraj, N., Kaibara, N., and Kuo, M.T. (2002). Expression of heavy subunit of γ-glutamylcysteine synthetase (γ-GCSh) in human colorectal carcinoma. International journal of cancer 97, 21–27.

51. Timmerman, L.A., Holton, T., Yuneva, M., Louie, R.J., Padró, M., Daemen, A., Hu, M., Chan, D.A., Ethier, S.P., and van’t Veer, L.J. (2013). Glutamine sensitivity analysis identifies the xCT antiporter as a common triple-negative breast tumor therapeutic target. Cancer cell 24, 450–465.

52. Ventura, A., Kirsch, D.G., McLaughlin, M.E., Tuveson, D.A., Grimm, J., Lintault, L., Newman, J., Reczek, E.E., Weissleder, R., and Jacks, T. (2007). Restoration of p53 function leads to tumour regression in vivo. Nature 445, 661–665.

53. Vyas, S., Zaganjor, E., and Haigis, M.C. (2016). Mitochondria and Cancer. Cell 166, 555–566.

54. Weinstein, C., Haschemeyer, R., and Griffith, O. (1988). In vivo studies of cysteine metabolism. Use of D-cysteinesulfinate, a novel cysteinesulfinate decarboxylase inhibitor, to probe taurine and pyruvate synthesis. Journal of Biological Chemistry 263, 16568–16579.

55. Winterbourn, C.C., and Hampton, M.B. (2008). Thiol chemistry and specificity in redox signaling. Free Radical Biology and Medicine 45, 549–561.

56. Yang, W.S., SriRamaratnam, R., Welsch, M.E., Shimada, K., Skouta, R., Viswanathan, V.S., Cheah, J.H., Clemons, P.A., Shamji, A.F., and Clish, C.B. (2014). Regulation of ferroptotic cancer cell death by GPX4. Cell 156, 317–331.

57. Zhang, Y., Tan, H., Daniels, J.D., Zandkarimi, F., Liu, H., Brown, L.M., Uchida, K., O’Connor, O.A., and Stockwell, B.R. (2019). Imidazole ketone erastin induces ferroptosis and slows tumor growth in a mouse lymphoma model. Cell chemical biology 26, 623–633. e629.

